# Structural and biochemical analysis of the Estrogen-Related Receptor α and complex with TMPRSS2 promoter DNA

**DOI:** 10.64898/2026.08.19.744156

**Authors:** K. Chandrashekhara, Ajay K. Saxena

## Abstract

In TMPRSS2 fusion-positive prostate cancer, ERRα is involved in regulation of ERG and promotes the androgen receptor independent signaling in the cancer progression. The ERRα binds to the ERREs (estrogen-related receptor response elements) present at −5042 bp of the TMPRSS2-promoter and enhances the ERG overexpression that causes prostate cancer progression. To dissect the structural basis of the ERRα recognition to the TMPRSS2 promoter DNA, we have purified the full-length ERRα (ERRαFL), NTD deleted construct (ERRαΔNTD), and the DNA-binding domain (ERRαDBD) proteins and performed the binding analysis with 30 bp TMPRSS2-promoter DNA (5′ -AGTCCAAGGTCGGTGGATCACAAGGTCAGG-3′). Circular dichroism analysis showed that all three ERRα proteins adopt native secondary structures. DNA binding induced subtle changes in the secondary structures, while enhancing the thermal stability (Tm) of all ERRα proteins. Binding analysis showed that ERRαDBD bound weakly to the DNA, whereas ERRαFL and ERRαΔNTD exhibited substantially higher affinities ∼120-fold and ∼131-fold than ERRαDBD, respectively. Small-angle X-ray scattering (SAXS) analyses revealed a dimeric ERRαFL structure and an ERRαFL-DNA complex (2:1) structure in solution and fitted well with Alpha Fold model of apo and DNA bound complex of ERRαFL. Furthermore, 100 ns dynamics simulations on apo and DNA-bound ERRα proteins showed that all proteins remained structurally stable, with flexibility largely confined to loop regions of ERRα proteins. Our biophysical, DNA binding and structural analyses have revealed the mechanism involved in ERRα recognition of the TMPRSS2-promoter DNA, which provides insight into ERRα-mediated transcriptional regulation and development of anticancer drugs against ERRα-driven prostate cancer.

## Introduction

ERRα (Estrogen-related receptor alpha) is an orphan nuclear receptor that exhibits constitutive transcriptional activity and independent of ligand binding [1]. The ERRα is involved in metabolic reprogramming and oncogenic progression in the colon, breast, ovarian and prostate, offering an attractive anticancer drug target [2,3]. The Activation function-2 (AF-2) of helix H12 in the ligand binding domain of ERRα interacts with coactivator Peroxisome proliferator-activated receptor gamma coactivator 1α (PGC-1α), stabilizes the active state and facilitating recruitment of coactivators [4]. The ERRα-PGC-1α complex regulates the genes involved in tricarboxylic acid cycle, oxidative phosphorylation, fatty acid β-oxidation, and lipid metabolism, which are essential for energetic and biosynthetic requirements of the cell [5].

The ERRα belongs to the nuclear receptor subfamily-3 and shares high homology to Estrogen-receptor alpha (ERα) in its DNA binding domain and does not require Estrogen for its activation [6,7]. The ERRα consists of four functional domains, (i) N-terminal transactivation domain (NTD) contains activation function-1 (AF-1) region, involved in ligand-independent activation, (ii) DNA-binding domain (DBD) with two zinc-finger motifs, (iii) Hinge region (HR) involved in nuclear localization and protein-protein interaction and (iv) Ligand binding domain (LBD) mediates homodimerization and contains the AF-2 pocket, which forms the coactivator-binding surface involved in recruitment of transcriptional coactivators [8].

The ERRα, Hepatocyte nuclear factor 4α (HNF-4α), and Peroxisome proliferator-activated receptor gamma (PPARγ) transcription factors belong to the NR3B1 nuclear receptor superfamily having similar structural fold and partially overlapping metabolic functions [9]. The crystal structures of the several receptors PPARγ-RXRα [10], RARβ-RXRα [11], LXRβ-RXRα [12] and HNF-4α [13] have been determined. These proteins form heterodimer and homodimer upon their response element, a direct 5’-AGGTCA-3’ repeat spaced by nucleotides respectively. In addition, ligand binding domain of the ERRα in complex with co-activator peptide from PGC-1α has been determined [14].

The ERRα promotes the tumor-stromal interactions that drive metabolic reprogramming and metastatic potential [2]. The ERRα is involved in unique metabolic profile of prostate tumors and regulate the citrate metabolism and zinc homeostasis. Several synthetic inhibitors i.e., (i) XCT790, inhibit the ERRα [15], (ii) Antagonist compound 29 [16] and (iii) indirect inhibitors disrupting the ERRα-PGC-1α interactions [17] have been identified and characterized.

The TMPRSS2-ERG (T-E) fusion gene is the most common genomic alteration in prostate cancer, transcriptionally regulated by ERRα through direct binding to ERRα response elements (ERREs) within the TMPRSS2 promoter [18, 19]. Three major ERREs have been identified at positions −635 bp (E1), −5,042 bp (E2), and −13,533 bp (E3) relative to the T-E fusion gene transcription start site. Among these, the E2 site (−5,042 bp) exhibits the strongest and most significant ERRα occupancy, indicating predominant functional ERRE mediating ERRα-dependent transcriptional activation of the TMPRSS2-ERG fusion gene [20]. In addition, regulatory interactions between ERRα and ERG promoter DNA in androgen receptor-independent prostate cancer progression, indicating ERRα as a promising therapeutic target.

In current study, we have used the integrative structural and biochemical approach to understand the molecular mechanism involved in ERRα and its TMPRSS2 promoter DNA recognition. We have used biochemical, small-angle X-ray scattering (SAXS) and molecular dynamics simulation techniques to understand the mechanism of sequence-specific DNA binding, domain organization, and conformational dynamics of ERRα during TMPRSS2 promoter DNA recognition. These data will provide insight into ERRα-mediated transcriptional regulation and will aid in rational design of therapeutic agents targeting ERRα-driven signalling in TMPRSS2-ERG fusion positive prostate cancer.

## Results

### Expression and purification of the ERRα proteins

A schematic diagram showing the bidirectional regulatory loop between ERG and ERRα in prostate cancer (Fig. 1A). The ERRα binds the ERRα-responsive elements (ERREs) at −5042 in TMPRSS2-ERG promoter region and activate the ERG expression. In turn, ERG binds to the ERRα promoter region at −605 and enhances the ERRα (transcription start site at +1). This entire positive feedback loop contributes to prostate cancer progression. A schematic view of full-length ERRαFL (1-423 amino acids), ERRαΔNTD (76-423 amino acids) and ERRαDBD (76-152 amino acids) where the ERRα consists of N-terminal transactivation domain (NTD, cyan), DNA-binding domain (DBD, red), Hinge region (HR, yellow), Ligand-binding domain (LBD, purple) and Activation function-2 region (AF-2, blue) (Fig. 1B). Alpha-fold modelled structure of the ERRα showing overall organization of the NTD, DBD, HR and LBD domains of the ERRα. NTD and HR regions of the ERRα form highly flexible loops, while DBD and LBD domains are highly structured (Fig. 1C). The topology diagram of ERRα shows the N-terminal domain (NTD), C4 zinc-finger motifs, α1 and α2 helices, an antiparallel β1-β2 sheet, and a 3₁₀ helix within the DNA-binding domain (DBD). The hinge region (HR) connects the DBD to the ligand-binding domain (LBD), which consists of helices α3-α12, with helix α12 forming the AF-2 coactivator-binding site (Fig. 1D).

**Fig. 1:**
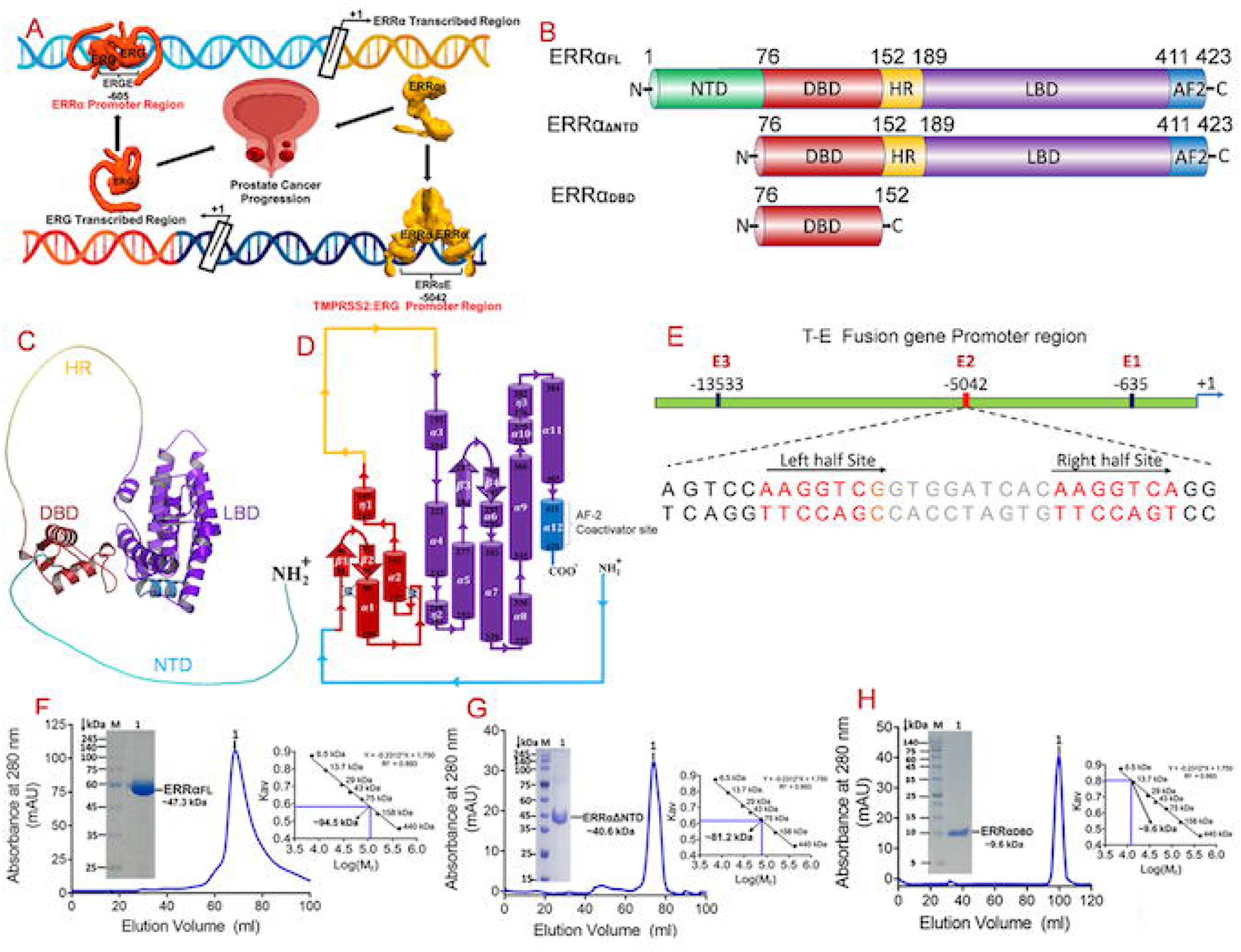
Schematic diagram of reciprocal transactivation between ERRα and ERG and Domain architecture, Alpha fold model of ERRαFL, structural topology, DNA recognition, Size-exclusion chromatography profiles of ERRα proteins. (A) Schematic model illustrating reciprocal transcriptional regulation between ERG and ERRα in prostate cancer. ERG binds the ERRα promoter (−605), enhancing ERRα transcription (+1). ERRα, in turn, occupies ERRαEs within the TMPRSS2: ERG promoter region (−5042), promoting ERG expression and contributing to prostate cancer progression. (B) Domain architecture of full-length human ERRα (ERRαFL, residues 1–423) and truncation constructs. ERRαFL comprises the N-terminal domain (NTD, 1–76), DNA-binding domain (DBD, 76–152), hinge region (HR, 152–189), ligand-binding domain (LBD, 189–423) and Activation function-2 (AF-2, blue). ERRαΔNTD lacks residues 1–76, and ERRαDBD corresponds to the isolated DBD (76–152). Residue boundaries are indicated. (C) Domain-integrated Alpha fold model of ERRαFL illustrating spatial coupling between the intrinsically disordered NTD (cyan), DBD (red) with Zn^2+^ ion, HR (yellow), LBD (purple), and AF-2 (blue) providing a structural framework for DNA recognition and transcriptional activation. (D) Topology diagram of ERRαFL showing the N-terminal domain NTD (cyan), DNA-binding domain (DBD, red) with C4 zinc-finger motifs (α1–α2), the hinge region (HR, yellow), the ligand-binding domain (LBD, purple) comprising helices α3–α11 and β3-b4 strand and Activation function-2 (AF-2, blue) comprising helix α12 forms the AF-2 coactivator interface. N-and C-termini residue boundaries are indicated. (E) Schematic diagram shows the ERRα-binding sites, (E1, E2, E3) located in the T-E fusion promoter. DNA Sequence of the E2 site representing two ERRα binding core Sequence AAGGTCA (red) between spacer DNA (gray). (F-H) Size-exclusion chromatography profiles from Superdex 200 (16/60) showing absorbance at 280 nm (mAU) versus elution volume (ml) for purified (F) ERRαFL (G) ERRαΔNTD and (H) ERRαDBD. Each panel includes an inset displaying a Coomassie-stained SDS-PAGE gel (M: molecular weight marker, 1: purified protein band) confirming the purity and apparent molecular weight of the respective protein construct. Apparent molecular masses of ERRαFL, ERRαΔNTD and ERRαDBD ∼94.5 kDa, ∼47.3 kDa and ∼9.6 kDa respectively were estimated from the standard Kav versus Log (Mr) calibration curve using molecular weight standards.

ERRα response element E2 in the TMPRSS2 promoter DNA identified two ERRE-like half-sites arranged as direct repeats. The left half-site (5′-AAGGTCG-3′) differs from the canonical ERRE core sequence (AAGGTCA) by a single nucleotide substitution at the seventh position. The right half-site AAGGTCA perfectly matches with the consensus sequence. These two half-sites are separated by a 9 bp spacer, forming a direct repeat (DR9)-like configuration, that may facilitate cooperative ERRα binding (Fig. 1E). This arrangement promotes cooperative binding of the ERRα homodimer to TMPRSS2-DNA and enhances the DNA-binding affinity and transcriptional activation of the TMPRSS2-ERG fusion gene.

The ERRαFL (Fig. 1F), ERRαΔNTD (Fig. 1G) ERRαDBD (Fig. 1H) were purified and eluted from Superdex200(16/60) column with apparent molecular masses of ∼94.5 kDa, ∼81.2 kDa and ∼9.6 kDa respectively, as estimated from the standard Kav versus Log (Mr) calibration curve using molecular weight standards. These elution profiles indicate that ERRαFL and ERRαΔNTD proteins exist predominantly as dimers in solution. In contrast, the ERRαDBD exist in monomeric state. SDS-PAGE analysis showed a single band for each purified protein, confirming its high purity.

### Secondary structure and Thermal denaturation analysis on ERRα proteins

Far-UV circular dichroism (CD) data of ERRαFL, ERRαΔNTD, and ERRαDBD proteins, and corresponding DNA-bound complexes, were collected in wavelength range of 200-250 nm and calculated the mean residue ellipticity (Δε, deg·dmol^-1^·cm^-1^). Secondary structure analysis showed α-helix/β-sheet contents of 42.8%/3.7% for ERRαFL, 50.7%/4.1% for ERRαΔNTD, and 32.8%/8.3% for ERRαDBD. These data were quite consistent to secondary structure composition of three ERRα proteins, indicating native conformations in the solution. Secondary structural analysis on three ERRα proteins in complex with DNA revealed α-helix/β-sheet contents of 45.9%/4.3% for ERRαFL-DNA, 52.8%/3.9% for ERRαΔNTD-DNA and 34.0%/9.2% for ERRαDBD-DNA (Fig. 2A-C).

**Fig. 2:**
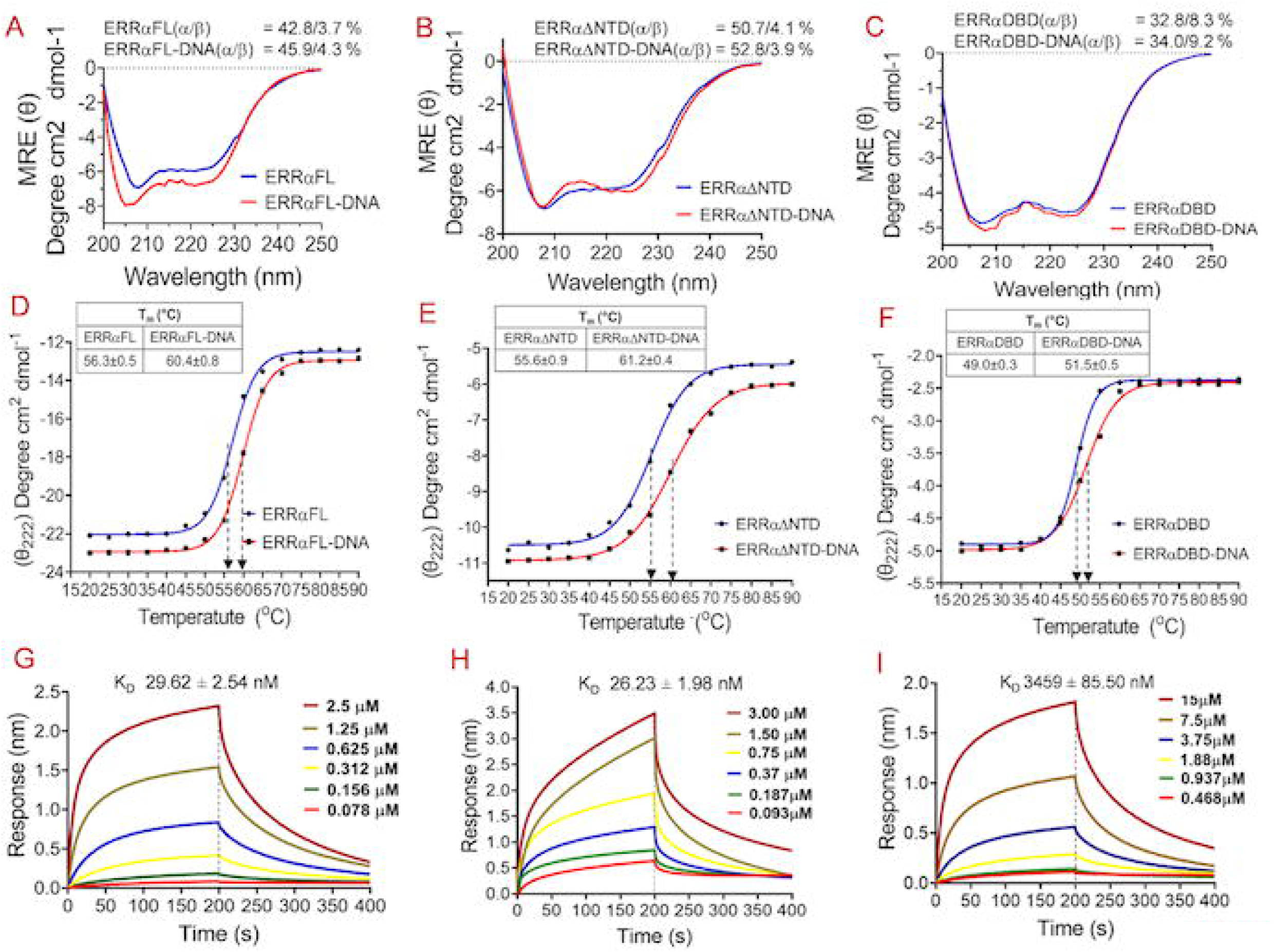
Secondary structure, thermal denaturation and DNA-binding profiles of the ERRα proteins. Circular dichroism spectra presenting the mean residue ellipticity (Δε mrw in M^-1^cm^-1^) as a function of wavelength (nm) for (A) ERRαFL (blue) and ERRαFL-DNA complex (red) (B) ERRαΔNTD (blue) and ERRαΔNTD -DNA complex (red) (C) ERRαDBD (blue) and ERRαDBD -DNA complex (red). The characteristic minima observed in the spectra indicate the presence of significant alpha-helical secondary structure within these ERRα proteins. For Thermal denaturation, mean residue ellipticity at θ222nm (Δεmrw in M^-1^cm^-1^) as a function of temperature (°C) for (D) ERRαFL (blue) and ERRαFL-DNA (red) (E) ERRαΔNTD (blue) and ERRαΔNTD -DNA (red) (F) ERRαDBD (blue) and ERRαDBD-DNA (red). Bio-layer interferometry (BLI) analysis demonstrating ERRα proteins binding to DNA. Sensorgrams showing the association and dissociation of (G) ERRαFL-DNA (H) ERRαΔNTD -DNA and (I) ERRαDBD -DNA. Global fitting of the kinetic data yielded a dissociation constant (K_D_) for ERRαFL∼29.62±2.54, ERRαΔNTD∼26.23±1.98 nM and ERRαDBD∼3459±85 nM.

Comparison of the secondary structures of the apo and DNA-bound three ERRα proteins indicate that DNA binding induced only subtle changes in the overall secondary structures of the proteins. Among them, ERRαΔNTD exhibited the smallest variation upon DNA binding, whereas ERRαFL showed a modest increase in both α-helical and β-sheet contents. Overall, data showed that DNA binding does not substantially alter the global secondary structure of three ERRα proteins. The DNA binding preserves the native fold of the three ERRα proteins, while accommodating local conformational adjustments associated with complex formation with DNA.

Thermal denaturation analysis of the ERRαFL, ERRαΔNTD and ERRαDBD, as well as their respective DNA-bound complexes, were performed after measuring CD signal at 222 nm, in 20-90 °C range 5°C intervals (Fig. 2D-F). The melting temperature (T_m_) of the apo ERRαFL was 56.3± 0.5 °C, which increased to 60.4 ± 0.8°C upon DNA binding. Similarly, ERRαΔNTD exhibited a T_m_ of 55.6 ± 0.9 °C in the unbound state, which increased to 61.2± 0.4 °C in the DNA-bound form. For ERRαDBD, the Tm was 49.0 ± 0.3 °C, which increased to 51.5 ± 0.5 °C upon DNA binding. Overall, these data indicated that DNA binding consistently increased the thermal stability of ERRα proteins, as reflected by the higher melting temperatures of the protein-DNA complexes compared their apo forms. The largest stabilization was observed for ERRαΔNTD, where Tm increased to ∼ 5.6 °C, compared with increases of 4.0 °C and 2.5 °C for ERRαFL and ERRαDBD respectively. These data indicated that DNA binding enhanced the stability of the folded structures of ERRα proteins, likely by strengthening protein-DNA interactions and reducing conformational flexibility. This interpretation is consistent with the accompanying CD secondary structure analysis, which showed modest changes in the secondary structural content of the ERRα-DNA complexes.

### Binding analysis of the ERRα proteins with TMPRSS2 promoter DNA

DNA-binding affinities of the ERRαFL, ERRαΔNTD and ERRαDBD proteins were determined following deletion of the respective functional domains of ERRα (Fig. 2G-I). The ERRαDBD exhibited very weak binding with DNA, having an equilibrium dissociation constant (*K_D_*) of ∼3459.00± 85.50 nM. In contrast, ERRαFL binds DNA with significantly higher affinity (*K_D_* ∼29.62± 2.54 nM), representing ∼117-fold increase in binding affinity.

These results indicated that ligand-binding domain of ERRα (LBD, residues 189-423) plays a critical structural role beyond ligand binding and substantially enhancing the DNA-binding affinity. The ERRαDBD alone is insufficient for stable and high-affinity DNA binding. The ERRαΔNTD binds DNA with little higher affinity (*K_D_* ∼26.23± 1.98) than ERRαFL, however significantly higher than ERRαDBD alone (∼131 fold). These data indicated that LBD and DBD modules of the ERRα are functionally integrated to establish the DNA binding affinity. It also indicated that ERRαNTD (1-76 amino acids) and ERRαHR (152-189 amino acids) domains are highly flexible and do not contribute significantly to the overall affinity of the ERRα to DNA.

### Low-resolution structures of the ERRαFL and ERRαFL-DNA complex using small angle X-ray scattering

The SAXS data on ERRαFL, and ERRαFL-DNA complex were measured at wavelength of 1.5418 Å at 283 K and converted into scattering intensity (I) ∼ angle (q) (Fig. 3A), which reflected the shape and size of the ERRαFL in solution. Details of the SAXS data collection and low-resolution structure analysis of ERRαFL and ERRαFL-DNA complex are given in Table 1. Radius of gyration 5.24 ± 0.4 nm from Guinier plot (Fig. 3B) and 5.25 ± 0.21 nm from P(R) curve (Fig. 3D) were observed for ERRαFL and showed good correlation to each other. These data showed ERRαFL dimer in solution, consistent to dimer observed in size exclusion chromatography. Ten independent *ab initio* simulations were performed without any symmetry restrictions and 252 dummy atoms were used to build the *de novo* envelope of ERRαFL (Fig. 3E-F). Maximum particle dimension ∼ 15.20±0.20 nm was observed for ERRαFL dimer and P(R) distribution analysis, showed consistent and symmetrical distribution, indicating a multisubunit symmetrical complex (Fig. 3D). Molecular weight analysis using forward scattering I(0) analysis (Fig. 3A) showed a Mw∼ 90.8 kDa, indicative of dimeric ERRαFL in the SAXS envelope (Fig. 3E-F). The dimeric ERRαFL model was well accommodated within the experimental SAXS envelope and showed good agreement with the experimental scattering curve. Flexible docking was performed on ERRαFL dimer using GROMACS program, as modelled ERRαFL dimer has little different orientations from dimeric ERRα-LBD structure observed in ERRα-PGC1α complex structure (PDB-3D24). The observed ERRαFL dimer fitted very well into low resolution SAXS envelope having good χ^2^ ∼2.43 (Table1, Fig. 3E-H).

**Fig. 3:**
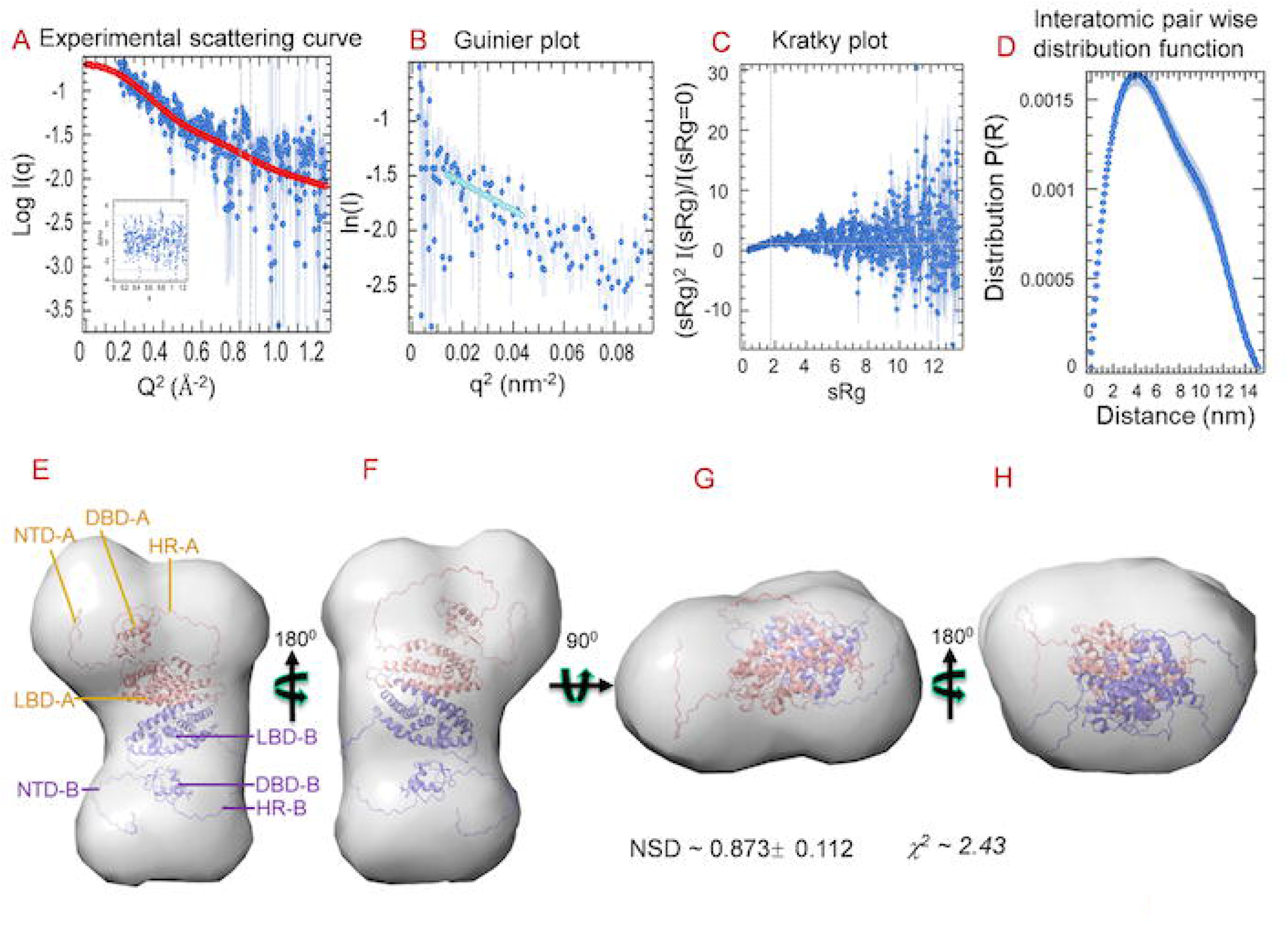
Low resolution structures of ERRαFL using small-angle X-ray scattering. (A) Experimental SAXS intensity profile of ERRαFL. The experimental data are shown as blue circles, and the fitted curve is represented by a red line. The inset shows the residuals of the fit, expressed as the ratio of intensity to its standard deviation plotted against s. (B) Guinier Plot of the natural logarithm of the scattering intensity versus the square of the scattering vector. The linear region at low s values (cyan) was used to determine the radius of gyration Rg. (C) Kratky plot in low q region showing the linear agreement between experimental data (blue circle) and Guinier equation (blue line). (D) Inter-atomic pairwise distribution function [P(r)] obtained from GNOM program. It shows the frequency distributions of interatomic vectors from predominant scattering species in the solution. At D_max_ ∼ 15.20 nM, the P(r) function approached zero smoothly. (E-F) 180^0^ view of the fitting of ERRαFL dimer [ERRa-A (orange) and ERRa-B (slate)] into experimental SAXS envelope generated by DAMFILT. (G-H) 180^0^ view of the fitting of ERRαFL dimer [ERRa-A (orange) and ERRa-B (slate)] into experimental SAXS envelope generated by DAMFILT. *De novo* envelope was obtained by superposing 10 models, averaged, and filtered by the DAMFILT program. Average NSD ∼ 0.873±0.112 was observed from ten aligned *de novo* models (n=10) and reported as (mean ± SD). The *de novo* models were quite similar, and resolution of ensemble was found ∼ 12.4 - 9.0 nm.

**Table 1.**
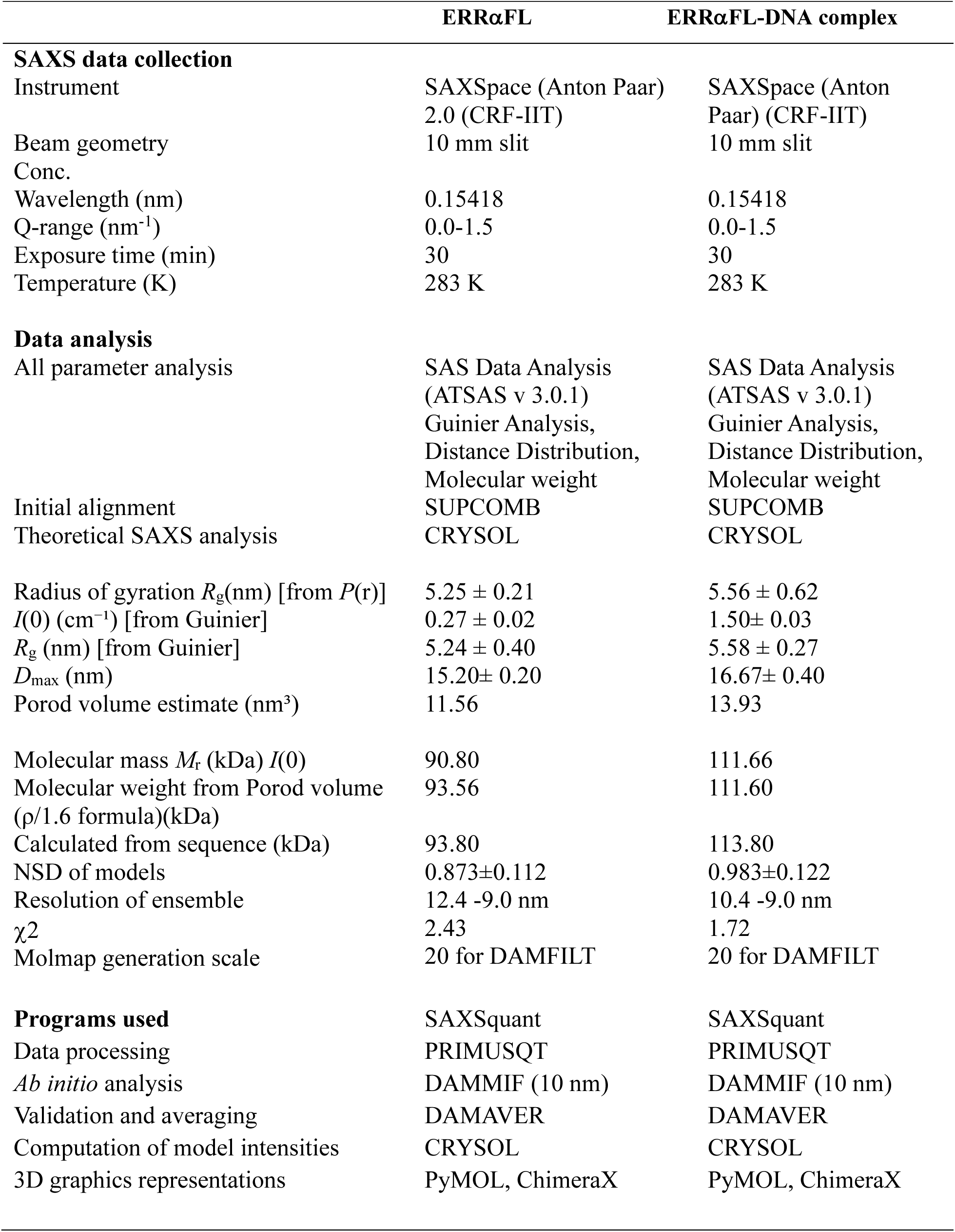
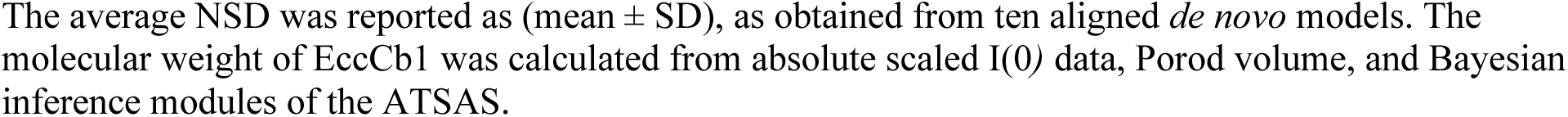
SAXS parameters used the low-resolution structure analysis ERRαFL and ERRαFL-DNA complex.

The SAXS data on ERRαFL-DNA complex was collected at 1.5418 Å in 283 K and converted into scattering intensity (I)∼ scattering angle (q) (Fig. 4A). Radius of gyration of 5.58 ± 0.27 nm from Guinier plot (Fig. 4B) and ∼ 5.56 ± 0.62 nm from P(R) (Fig. 4D) curve were observed for ERRαFL-DNA complex. Maximum particle dimension ∼16.67± 0.40 nm was observed from P(R) distribution analysis (Fig. 4D) which showed consistent and symmetrical distribution, indicative of 2:1 stoichiometry ERRαFL-DNA in solution. Molecular weight analysis using forward scattering I(0) indicated Mw ∼111.6 kDa, which corresponds to (2:1) complex of ERRαFL-DNA (2:1 stoichiometry in solution (Fig. 4E-H). 611 dummy atoms were used to develop the low-resolution envelope of the ERRαFL-DNA complex and fitted very well with Alpha fold modelled ERRαFL-DNA (2:1) complex which has χ^2^ ∼1.72 (Table 1, Fig. 4E-H).

**Fig. 4:**
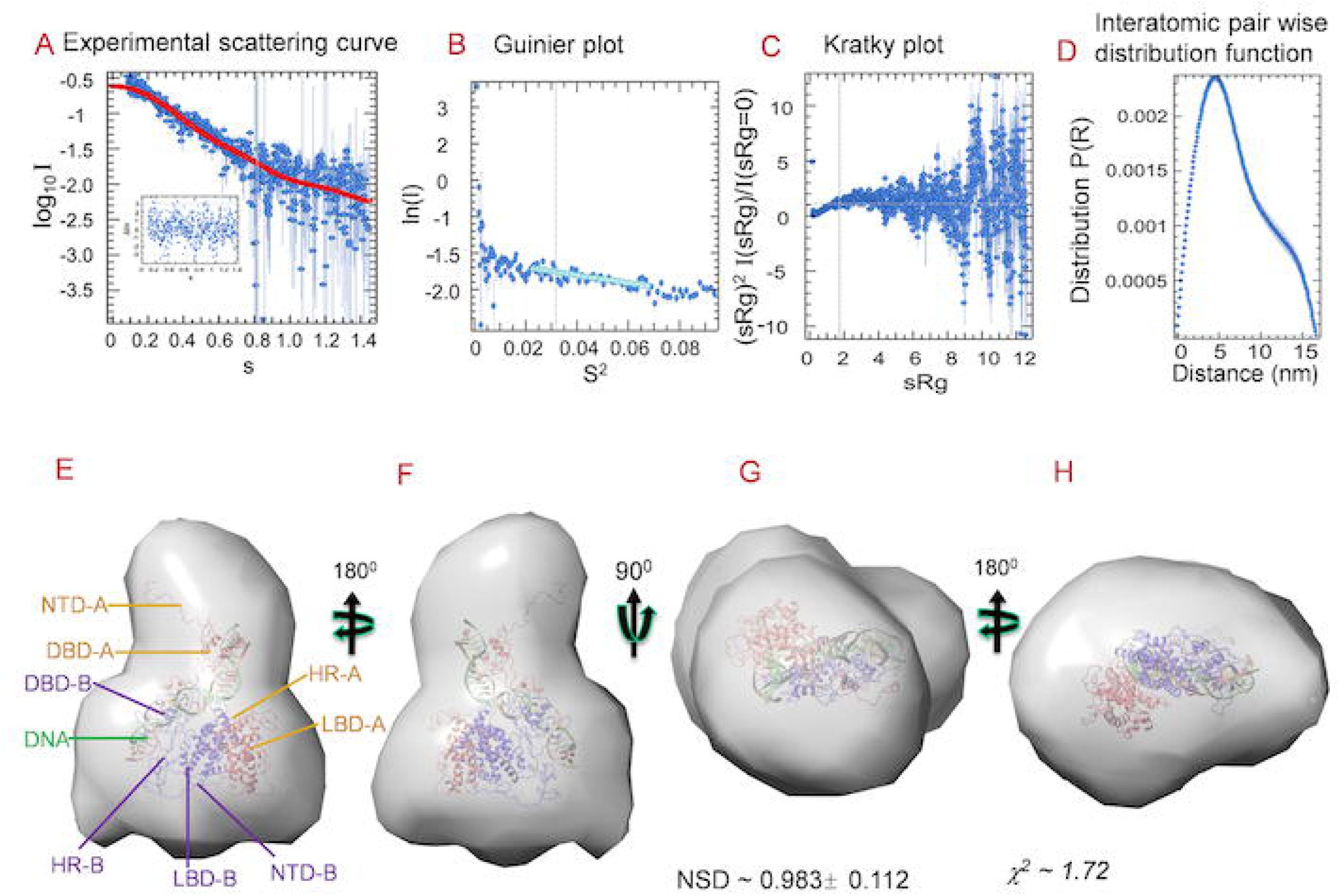
Low resolution structures of ERRαFL-DNA complex using small-angle X-ray scattering. (A) Experimental SAXS intensity profile of ERRαFL-DNA complex. The experimental data are shown as blue circles, and the fitted curve is represented by a red line. The inset shows the residuals of the fit, expressed as the ratio of intensity to its standard deviation plotted against s. (B) Guinier Plot of the natural logarithm of the scattering intensity versus the square of the scattering vector. The linear region at low s values (cyan) was used to determine the radius of gyration Rg. (C) Kratky plot in low q region showing the linear agreement between experimental data (blue circle) and Guinier equation (blue line). (D) Inter-atomic pairwise distribution function [P(r)] obtained from GNOM program. It shows the frequency distributions of interatomic vectors from predominant scattering species in the solution. At D_max_ ∼ 16.67 nM, the P(r) function approached zero smoothly. (E-F) 180^0^view of the fitting of ERRαFL-DNA complex [ERRa-A (orange), ERRa-B (slate) and DNA (green)] into experimental SAXS envelope generated by DAMFILT. (G-H) 180^0^ view of the fitting of ERRαFL-DNA complex [ERRa-A (orange), ERRa-B (slate) and DNA (green)] into experimental SAXS envelope generated by DAMFILT. *De novo* envelope was obtained by superposing 10 models, averaged, and filtered by the DAMFILT program. Average NSD ∼ 0.983±0.112 was observed from ten aligned *de novo* models (n=10) and reported as (mean ± SD). The *de novo* models were quite similar, and resolution of ensemble was found ∼ 10.4 - 9.0 nm.

### Overall structure of the ERRαFL dimer

Interface analysis on ERRαFL dimer (Fig. 5A) showed 1287 Å^2^ buried surface area (total ∼ 25967 Å^2^) between two monomers ERRα-A and ERRα-B, in which LBD-LBD domain shares 1380 Å^2^ buried surface area. The ERRαFL dimeric interface is mediated by α7, α8, α9, and α10 helices, with 12 hydrogen bonds and 8 salt bridges observed at the dimeric interface (Table S1). Quite similar interfaces were observed in the structures of HNF-4α-DR1 (PDB-4IQR), Glucorticoid receptor-DR4 (PDB-7PRW) and ERRα-LBD dimer (PDB-3D24). The hydrogen-bonding network includes interactions between Gln311(α7) and Asp338 (loop); Arg315 (α7) and Asp338 (loop)/Glu349 (α9); Asn336 (α8) and Arg315 (α7)/Leu383 (loop); Asp338 (loop) and Gln311 (α7)/Arg315 (α7); Glu353 (α9) and Arg380 (α10); Leu360 (α9) and Arg375 (η3); Arg375 (η3) and Leu360 (α9); Arg380 (α10) and Glu349 (α9); and Leu383 (loop) and Asn336 (α8). In addition, salt bridges between Arg315 and Asp338, Glu353 and Arg380; Glu357 and Arg376 and their corresponding reciprocal interactions were observed, further stabilize the LBD dimeric interface. A total of 156 non-bounded contacts are invalid between the ERRα-A and ERRα-B LBD interphase.

**Fig. 5:**
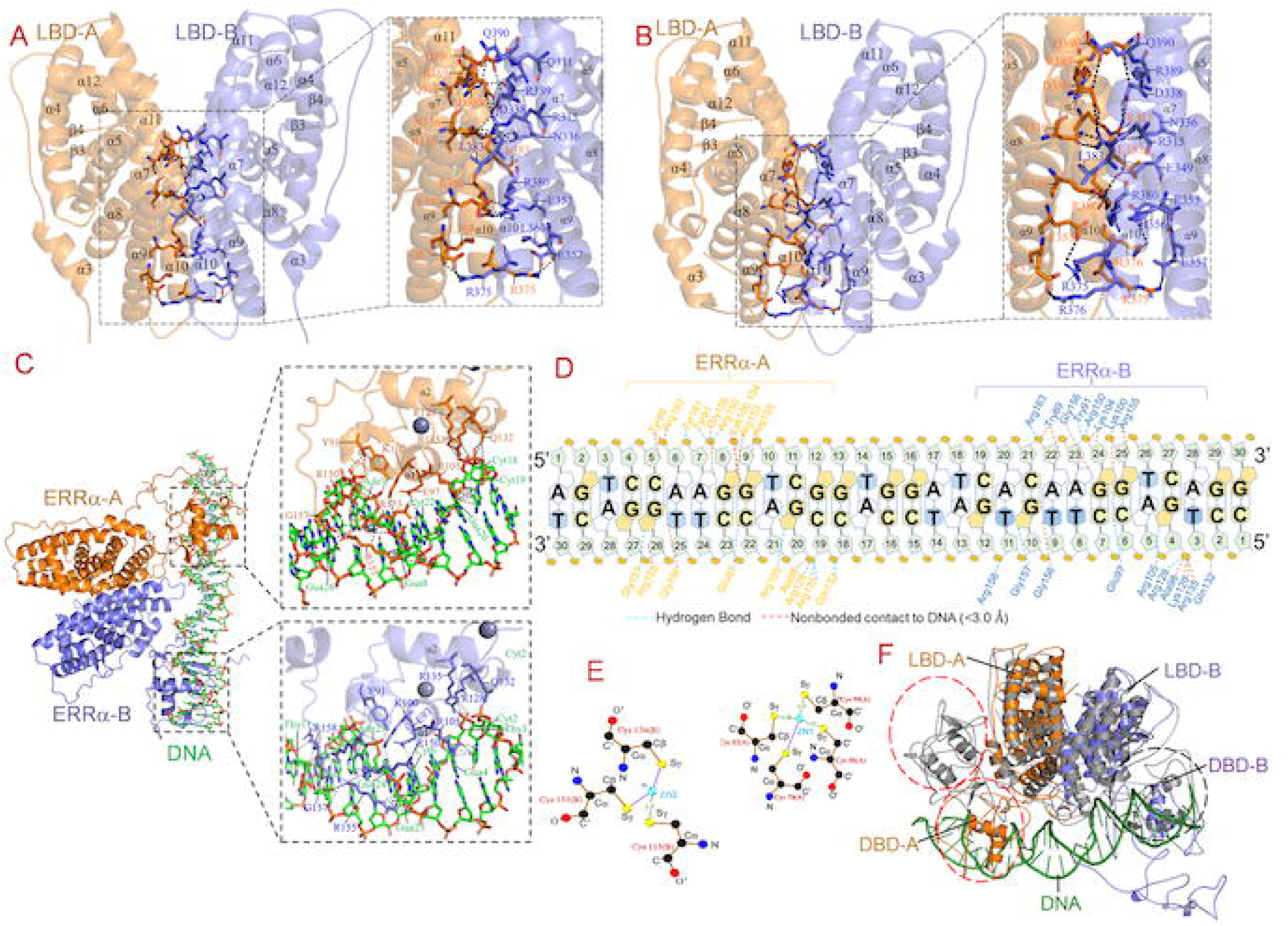
Overall structure of ERRα dimer and ERRα-DNA complex. (A) LBD–LBD interface in apo ERRα. Left: Overall three-dimensional structure of the ERRa dimer, with the two interacting ERRα-A (orange) and ERRα-B (slate) cartoon representations. Right: Enlarged view of the binding interface highlighting key interacting amino acid residues. Side chains of interface residues are displayed as sticks and labelled to illustrate the molecular interactions. (B)LBD–LBD interface in ERRα-DNA complex. Left: Overall three-dimensional structure of the ERRa-DNA complex, with the two interacting ERRα-A (orange) and ERRα-B (slate) cartoon representations. Right: Enlarged view of the binding interface highlighting key interacting amino acid residues. Side chains of interface residues are displayed as sticks and labelled to illustrate the molecular interactions. (C) Ribbon diagram of the ERRα-A (orange) and ERRα-B (slate) homodimer interaction with DNA (green). Closeup view of the hydrogen bonding interactions between DBD to ERRα-A (orange) and ERRα-B (slate). (D) Schematic diagram of the detailed interactions between ERRα-A (orange) and ERRα-B (slate) molecules on DNA. The hydrogen bonds and non-bonded contacts are shown in blue and orange dotted lines respectively. (E) Ligplot showing the co-ordination and hydrogen bonds between two Zn^2+^ ion with zinc-Figure motif residues of ERRαDBD. (F) Superposition of the ERRα dimer (grey) on ERRα dimer (orange and slate)-DNA (green) complex. DBD domains between both proteins showed major conformation while LBD domain was quite conserved.

### Overall structure of the ERRαFL-DNA complex

#### LBD-LBD interface

In the ERRαFL-DNA complex (Fig. 5B), the LBDs maintaining the head-to-head arrangement, while the DBDs engage the DNA in a head-to-tail orientation. 10 hydrogen bonds and 14 salt bridges were observed across the LBD dimer interface (Table S2), which stabilized the DNA-bound ERRαFL dimer. The hydrogen-bonding network includes interactions between Gln311(α7) and Asp338 (loop); Arg315 (α7) and Asn336 (α8)/Asp338 (loop); Asn336(α8) and Arg315(α7)/Leu383 (loop); Asp338 (loop) and Arg315 (α7); Glu349 (α9) and Arg380 (α10); Leu383 (loop) and Asn336 (α8); Glu353 (α9) and Arg375 (η3); Arg375 (η3) and Glu357 (α9); and Arg380 (α10) and Glu349 (α9). The salt bridges further reinforce the interface, including reciprocal Arg315 and Asp338 interactions, Glu349 and Arg380; Glu353/Glu357 with Arg376 and Arg380; Glu357 with Arg375 and Arg376, Glu361 with Arg375, reciprocal Arg375–Glu357/Glu361 interactions, Arg376 with Glu353/Glu357, and Arg380 with Glu349/Glu353. A total of 171 non-bounded contacts are invalid between the ERRα-A and ERRα-B LBD interphase. These data indicated that DNA binding strengthens the LBD dimeric interface, thereby contributing to stabilization of the ERRαFL-DNA (2:1 stoichiometry) complex.

#### DBD-DNA interface

In the ERRαFL-DNA complex, the DNA-binding domains of two monomers ERRα-A and ERRα-B bind tandemly two half-sites of DNA in a head-to-tail orientation (Fig. 5C). The α1 helix of the DNA-binding domain inserts into the DNA major groove, where it specifically recognizes 5′-CAAGGTCAG-3′ sequences through hydrogen bonding and van der Waals interactions. The DNA-binding domains of two monomers, ERRα-A and ERRα-B, bind complementarily to the DNA, burying surface areas of 303.7 and 284.1 Å^2^ respectively. In contrast, DBD-DNA interface in HNF4α-DR1 complex (PDB-4IQR) and glucocorticoid receptor-DR4 complex (PDB-7PRW) share buried surface areas of 303.7 and 284.1 Å^2^ respectively.

Detailed analysis of the ERRα-DNA interface revealed an extensive hydrogen bonding network (Fig. 5D). ERRα-A residues on the left half-site of DNA 5′→3′, Arg150 (loop) and Tyr91 (β-turn) form bifurcated hydrogen bonds with PO_4_ moiety of Ade7. On the 3′→5′site, Gln132(α2) forms a hydrogen bond with the phosphate group of Cyt18, Arg128(loop) and Arg135 (α2) form trifurcated hydrogen bonds with the phosphate group of Cyt19 and Arg105(α1), Glu97(α1) and Gly157(loop) form hydrogen bond with the nitrogenous bases of Gly20, Cyt22 and Gly26 respectively. ERRα-B residues on the right half-site of DNA 5′→3′, Arg150 (loop) and Tyr91 (β-turn) form bifurcated hydrogen bonds with PO_4_ group of Ade23. On the complementary 3′→5′site, Arg128(α2) forms a hydrogen bonds with the phosphate group of Tyr3, Arg105(α1), Glu97(α1) and Gly157(loop) form hydrogen bond with the nitrogenous bases of Gly4, Cyt6 and Gly10 respectively. Arg158 (loop) forms a hydrogen bond with the phosphate group of Tyr11.

The two Zinc atoms, the Zn1 forms coordination geometry with Sγ atoms of the Cys79, Cys82, Cys 96 and Cys99 residues, while Zn2 forms with Cys115, Cys118, Cys131, and Cys134 residues of the ERRα monomer (Fig. 5E).

Superposition of the ERRαFL dimer (grey) on the ERRαFL-DNA complex [ERRαA (orange), ERRαB (slate) and DNA (green)] yielded RMSD ∼7.3 Å for 572 Cα atoms (Fig. 5F). The structure of the LBDs was quite conserved between apo, and DNA bound ERRαFL, however orientations of DBDs changed significantly upon complex formation with DNA.

### Sequence alignment and comparative structure analysis

DALI server analysis identified proteins from the nuclear receptor superfamily as the closest structural homologs of ERRα. The glucocorticoid receptor exhibited a Z-score of ∼24.0, ∼27% sequence identity, and an RMSD of ∼4.5 Å across 237 aligned residues out of a total of 326 residues. Similarly, HNF4α displayed a Z-score of ∼20.1, ∼28% sequence identity, and an RMSD of ∼6.1 Å across 215 aligned residues out of 303 residues. These findings showed that ERRα adopts a conserved structural fold characteristic of the nuclear receptor family despite relatively low sequence identity.

The ERRα sequence was aligned with Glucocorticoid receptor (PDB-7PRW) and HNF4α (PDB-4IQR) followed by mapping of secondary structures onto the sequence alignment where the ERRαDBD residues are involved in DNA binding ($$), Zn^2+^ binding ($$), LBD residues involved in dimerization (#) and disease-associated residues in Triple-negative invasive lobular carcinomas (TN-ILCs) (*) (Fig. 6A). The residues involved in the DNA-binding and TN-ILCs disease were quite conserved across Glucocorticoid and HNF4-α receptors, highlighting the evolutionary conservation of DNA recognition and disease mechanism. In contrast, residues involved in LBD-LBD dimerization were less conserved, however retain highly similar LBD-LBD dimerization interface and conservation of the overall structural architecture.

**Fig. 6:**
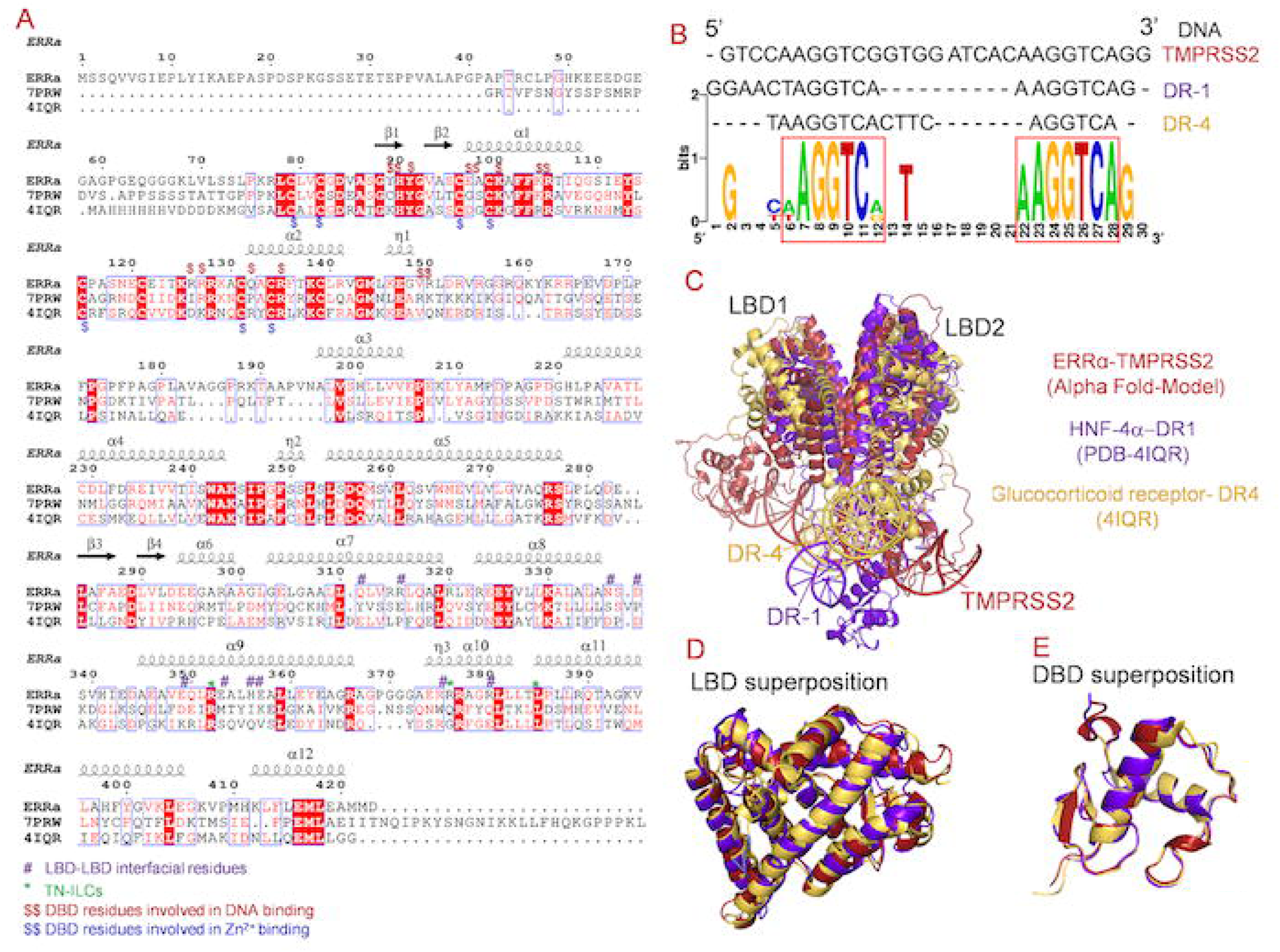
Sequence alignment, and comparative structural analysis of the ERRα. (A) Multiple sequence alignments of ERRαFL (1–423 residues) onto glucocorticoid receptor (DR2, PDB-7PRW) and hepatocyte nuclear factor 4α (DR4, PDB-4IQR) using MultiAln and ESPript3 program and mapped ERRαFL structure on sequence alignment. DNA-contacting residues within the DNA-binding domain (DBD) were marked ($, $), residues mediating LBD–LBD dimerization (#) and disease-associated mutations (*). (B) WebLogo Alignment of ERRα-responsive elements within the TMPRSS2 promoter showing direct repeat motifs with DR-1 and DR-4 response elements. Conserved AGGTCA half-sites are boxed; sequence logo (information content in bits) demonstrates high conservation of the core ERRE consensus. (C) The structure of the glucocorticoid receptor-DR4 complex (yellow, PDB-7PRW,) and HNF4α-DR1 complex (purple, PDB-4IQR) were superposed on ERRαΔNTD-TMPRSS2 complex (red, Alpha fold model) using the Superpose program. (D)LBD superposition of glucocorticoid receptor (yellow) and HNF4α (purple) were superposed on ERRα (red). (E) DBD superposition of glucocorticoid receptor (yellow) and HNF4α (purple) were superposed on ERRα (red).

**Fig. 7:**
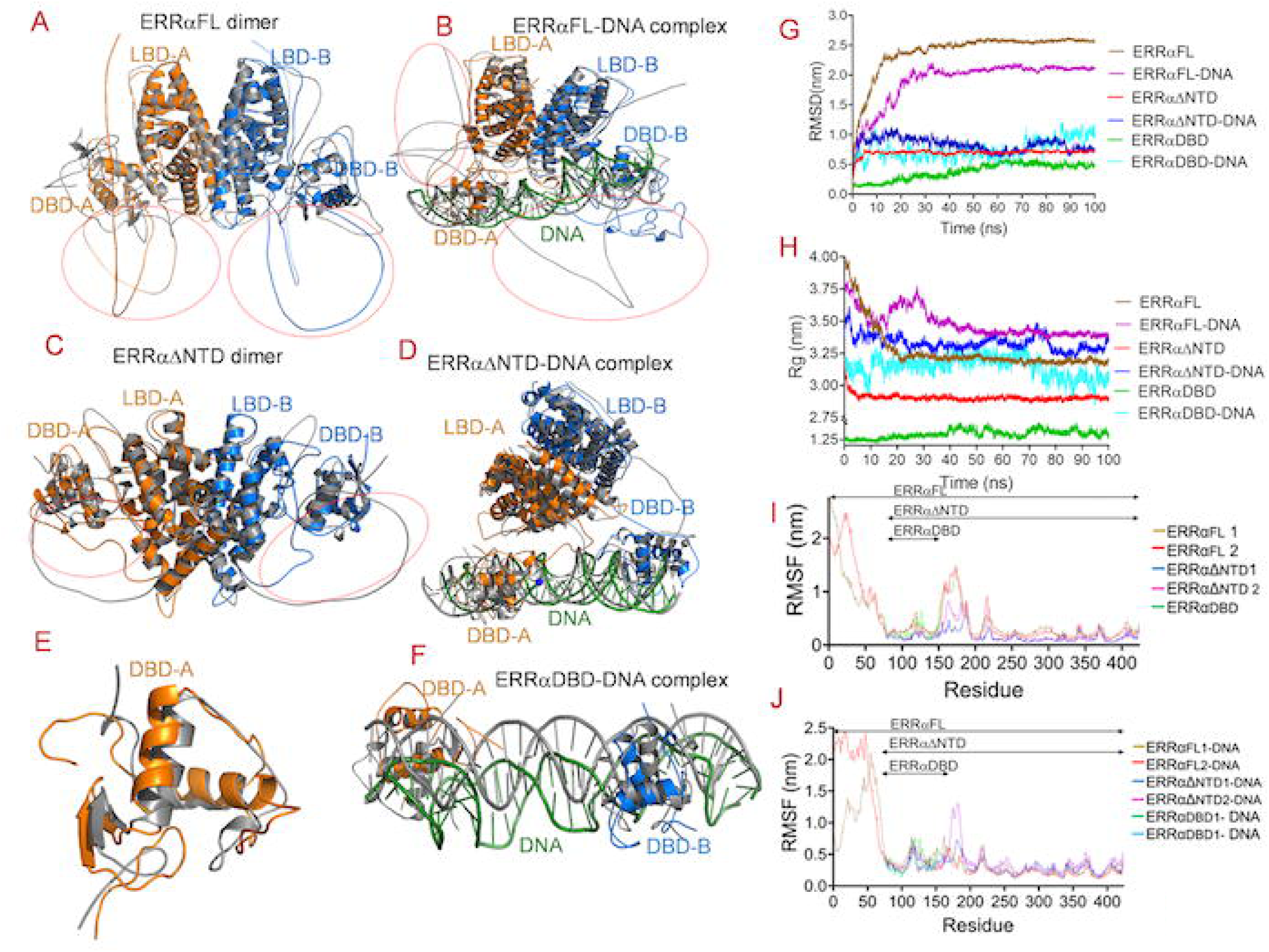
Molecular dynamics assessment of structural stability and compactness in ERRα variants and TMPRSS2 promoter DNA complexes. (A)Superposition of 100 ns simulated ERRαFL (orange and slate) on starting 0 ns ERRαFL (grey). (B)Superposition of 100 ns simulated ERRαFL-DNA complex (ERRαFL: orange and slate, DNA: green) on starting 0 ns ERRαFL-DNA complex (grey). (C) Superposition of 100 ns simulated ERRαΔNTD (orange and slate) on starting 0 ns ERRαΔNTD (grey). (D) Superposition of 100 ns simulated ERRαΔNTD -DNA complex (ERRαΔNTD: orange and slate, DNA: green) on starting 0 ns ERRαΔNTD -DNA complex (grey). (E)Superposition of 100 ns simulated ERRαDBD (orange) on starting 0 ns ERRαDBD (grey). (F)Superposition of 100 ns simulated ERRαDBD-DNA complex (ERRαDBD: orange and slate, DNA: green) on starting 0 ns ERRαΔNTD -DNA complex (grey). (G)RMSD Root-mean-square deviation profiles of Cα atoms over 100 ns all-atom MD simulations quantify time-resolved structural deviations and global compactness for full-ERRαFL, ERRαΔNTD, ERRαDBD and their DNA-bound complexes. (H) Radius of gyration (*Rg*) plot over 100 ns all-atom MD simulations quantify time-resolved structural deviations and global compactness for ERRαFL, ERRαΔNTD, ERRαDBD their DNA-bound complexes. (I) RMSF of over 100 ns all-atom MD simulations quantify time-resolved structural deviations and global compactness for ERRαFL, ERRαΔNTD and ERRαDBD. (J)RMSF of over 100 ns all-atom MD simulations quantify time-resolved structural deviations and global compactness for ERRαFL-DNA, ERRαΔNTD-DNA and ERRαDBD-DNA complexes

WebLogo diagram of the sequence alignment of the TMPRSS2-DNA (30 base pairs) on the DR1 of the HNF4α (PDB-4IQR) and DR4 of the Glucocorticoid receptor (PDB-7PRW) showed two consensus core 5’-AGGTCA-3’ and 5’-AAGGTCA-3’ site (Fig. 5B). Superposition of the full-length HNF4α-DR1 complex (cyan, PDB-4IQR) onto the ERRαFL-DNA complex (slate) yielded an overall RMSD of ∼5.53 Å over 441 Cα atoms, indicating substantial conservation of the overall structure (Fig. 6C). In contrast, superposition of the Glucocorticoid receptor (green, PDB −7PRW) onto the ERRαFL-DNA complex (slate) resulted in high RMSD of ∼22.71 Å over 527 Cα atoms. This data showed pronounced differences in the relative orientations of the DBD and DNA rather than divergence in their individual folds (Fig. 6C).

Domain-wise structural superposition of the ERRα showed high conservation of the LBD and DBD domains. Superposition of the HNF4α-LBD onto the ERRα-LBD yielded an RMSD of ∼1.52 Å over 174 Cα atoms (Fig. 6D), whereas the ERRα-DBD superimposed with an RMSD of ∼0.77 Å over 66 Cα atoms (Fig. 6E). Comparable RMSD values were obtained for the Glucocorticoid receptor. These data indicated that LBD and DBD domains of ERRα adopt highly similar fold, despite substantial differences in the arrangement of these domains within the full-length receptors.

Together, this result showed that ERRα preserves the canonical nuclear receptor fold and shares highly conserved DNA-binding and dimerization interfaces with Glucocorticoid and HNF4α nuclear receptors. The larger RMSDs in ERRαFL superpositions reflect the differences in interdomain organization and conformational flexibility, rather changes in the conserved core structures of the DBD and LBD. The strong conservation of DNA-contacting and disease-associated residues of ERRα further supports the common key functional mechanisms across nuclear receptor family.

### Dynamics simulation on apo and DNA bound ERRα proteins

100 ns molecular dynamics simulation was performed on ERRαFL, ERRαΔNTD, and ERRαDBD as well as their DNA bound complexes to investigate their conformational dynamics and structural stability throughout the simulation (Table 2). To assess conformational stability, simulated structures of apo and DNA bound ERRαFL, ERRαΔNTD, ERRαDBD complexes were superposed on starting structures and root-mean-square deviation (RMSD) were calculated for all Cα atoms. The average RMSD ∼1.8 Å for 544 Cα atoms were observed for ERRαFL (Fig. 6A) and ∼2.3 Å for 532 Cα atoms ERRαFL-DNA complex (Fig. 6B). ∼1.5 Å for 450 Cα atoms of ERRαΔNTD dimer (Fig. 6C) and ∼5.7 Å for 656 Cα atoms of ERRαΔNTD-DNA complex (Fig. 6C-D). ∼ 3.0 Å for 58 Cα atoms of ERRαDBD (Fig. 6A) and ∼4.4 Å for 86 Cα atoms of ERRαDBD-DNA complex (Fig. 6E-F).

**Table 2.** Secondary structures of ERRαFL, ERRαΔNTD, ERRαDBD and thermal stability profile.

| Proteins | Alpha-fold model |  |  | CD data |  |  | NRMSD | T<sub>m</sub> (°C) |
| --- | --- | --- | --- | --- | --- | --- | --- | --- |
| | α-helix | β-Sheet | Coil | α-helix | β-Sheet | coil | | |
| ERRαFL | 43.0 | 3.5 | 53.4 | 42.8 | 3.7 | 53.5 | 0.0262 | 56.3±0.5 |
| ERRαFL-DNA | 43.0 | 3.5 | 53.4 | 45.9 | 4.3 | 49.8 | 0.0543 | 60.4±0.8 |
| ERRα ΔNTD | 50.7 | 4.3 | 45.0 | 50.7 | 4.1 | 45.2 | 0.0342 | 55.6±0.9 |
| ERRα ΔNTD-DNA | 50.7 | 4.3 | 45.0 | 52.8 | 3.9 | 43.3 | 0.0611 | 61.2±0.4 |
| ERRαDBD | 32.0 | 7.7 | 60.3 | 32.8 | 8.3 | 58.9 | 0.0231 | 49.0±0.3 |
| ERRαDBD-DNA | 32.0 | 7.7 | 60.3 | 34.0 | 9.2 | 56.8 | 0.0228 | 51.5±0.5 |

**Table 3.**
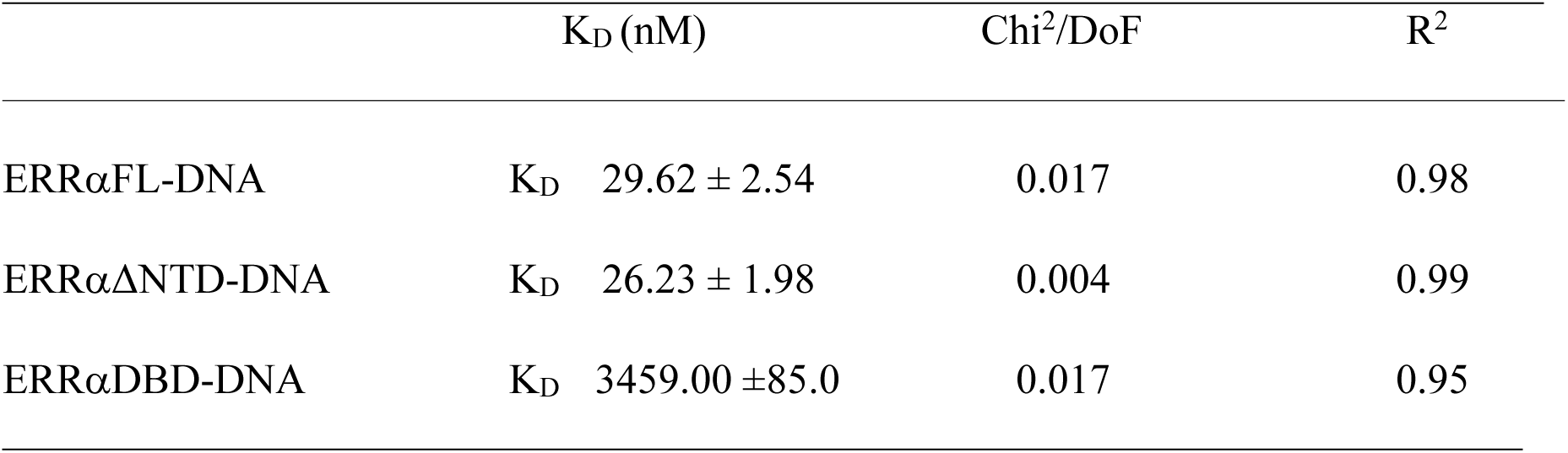
Binding analysis between ERRαFL, ERRαΔNTD, ERRαDBD with DNA.

**Table 4.**
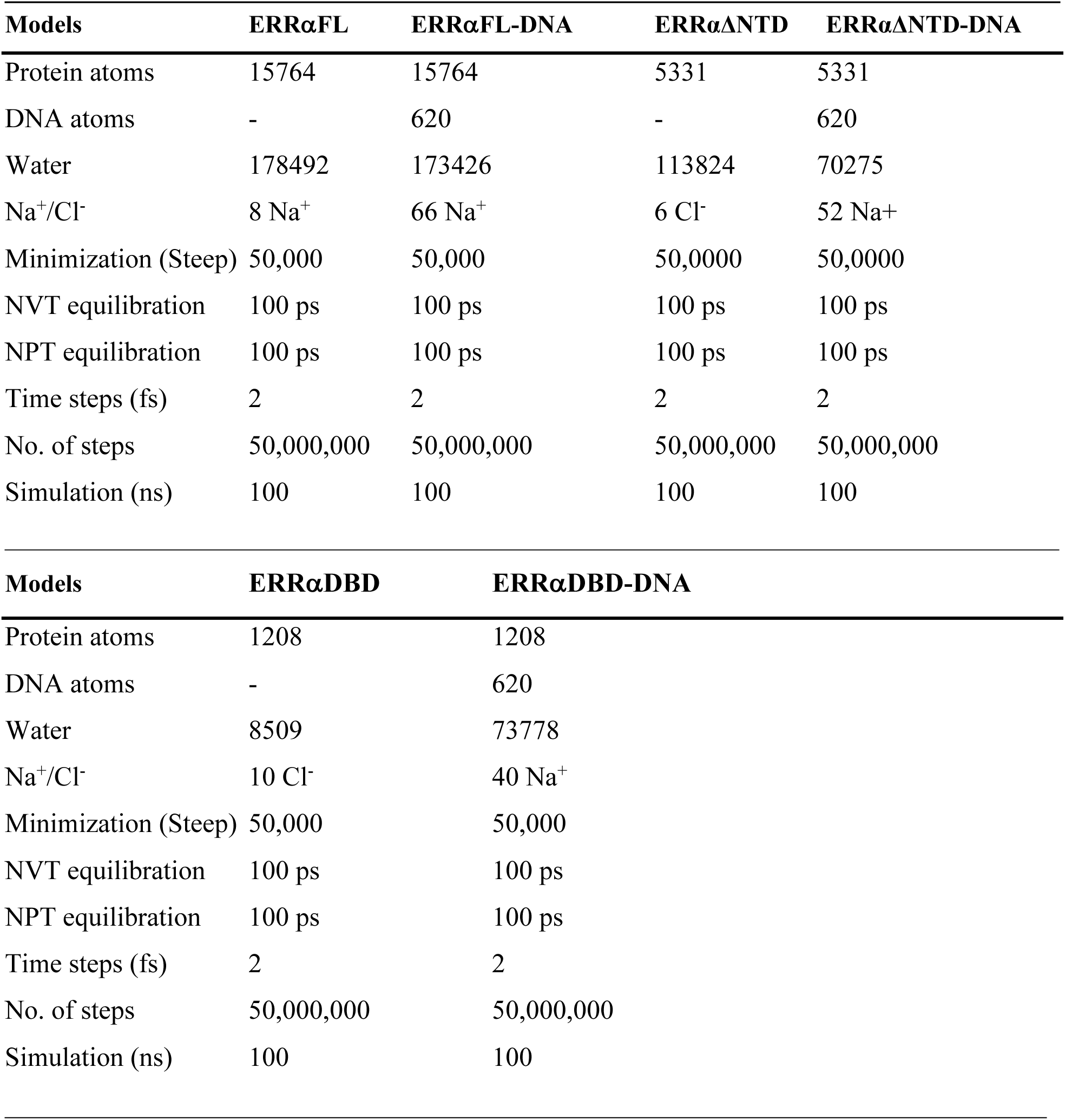
Various parameters used in dynamic simulations of ERRαFL, ERRαΔNTD, ERRαDBD and their complexes with DNA.

Overall, α-helices and β-strands in all ERRα in apo and DNA bound states quite conserved throughout 100 ns simulations. Higher flexibility was observed in HR loop regions of the apo, and DNA bound ERRα proteins. ERRαFL showed intermediate stability, reflecting contributions from both structured and flexible domains. ERRαΔNTD exhibited comparatively lower RMSD fluctuations, indicating greater structural stability relative to the ERRαFL. The ERRαDBD-DNA complex displayed higher conformational variability for protein and DNA.

Backbone RMSD trajectories for apo and DNA bound ERRαFL, ERRαΔNTD, and ERRαDBD were calculated as shown in Fig. 6G. The average RMSD values were ∼2.45 Å for ERRαFL, ∼ 2.18 Å for ERRαFL-DNA, ∼ 0.65 Å for ERRαDNTD, ∼ 0.68 Å for ERRαΔNTD-DNA, ∼ 0.42Å for ERRαDBD and ∼ 0.53 Å for ERRαDBD-DNA complex was observed. The ERRαΔNTD exhibited comparatively lower RMSD fluctuations than ERRαFL and ERRαDBD, which indicate greater structural stability relative to ERRαFL and ERRαDBD proteins.

Radius of gyration (Rg) was calculated to evaluate protein folding and structural compactness from the simulation trajectories (Fig. 6H). Following Rg values i.e., ∼32.5 Å for ERRαFL, ∼35.2Å for ERRαFL-DNA complex, ∼28.4 Å for ERRαΔNTD & ∼33.2 Å, for ERRαΔNTD-DNA complex and ∼ 12.5 Å for ERRαDBD & ∼ 31.5 Å for ERRαDBD-DNA complex were observed. ERRαΔNTD exhibited the most compact and stable conformation, whereas ERRαDBD showed greater expansion and flexibility. The Rg suggested that all proteins remained stable structurally with no indication of global unfolding.

Root-mean-square fluctuation (RMSF) was calculated to identify the localized structural perturbations during 100 ns simulation. In ERRαFL1 and ERRαFL2 (Fig. 6I), the RMSF goes from ∼25 −2 Å for 1-98 Cα atoms, ∼2-15 Å for 150-200 Cα atoms and remained stable ∼2 Å for other Cα atoms of both proteins. For ERRαΔNTD1, and ERRαΔNTD2, it remained stable ∼2 Å for Cα atoms of both proteins. For ERRαDBD, it remained stable to ∼2 Å for Cα atoms. Collectively, similar RMSF patterns were observed for the DNA bound ERRαFL, ERRαΔNTD and ERRαDBD proteins (Fig. 6J).

These data showed that backbone flexibility is predominantly localized in NTD region (1-90 residues) and 153-189 residue loop connecting to DBD to LBD domains (HR region) of the ERRα protein. The backbone Cα atoms of the LBD and DBD domains of ERRα protein remained constant during entire 100 ns simulation with no global conformational change. No significant conformational changes in different regions of ERRα undergo upon DNA binding, except minor fluctuations in NTD and HR loops regions of the protein.

#### Dynamics involved in hydrogen bonding

Both, inter and intra molecular hydrogen bonds dynamics within apo and DNA bound ERRαFL, ERRαΔNTD and ERRαDBD were calculated over 100 ns dynamics simulation. Hydrogen bonds were calculated across the trajectories to analyze the intramolecular stability. Hydrogen bonds consistently present, though number of intramolecular hydrogen bonds fluctuated during 100 ns simulation (Fig. S1A). Few hydrogen bonds exhibited relatively high frequency (∼46% and ∼28%, respectively) in occupancy analysis and maintained domain stability.

Intermolecular hydrogen bonds for apo and DNA bound ERRαFL, ERRαΔNTD, ERRαDBD were calculated to examine temporal and stability behaviour (Fig. S1B). These data showed stable and persistent hydrogen bonds with between ERRαFL, ERRαΔNTD, and within DNA. In ERRαDBD, frequent interactions with DNA were observed throughout 100 ns simulation. These data indicated that DNA establishes a stable binding network within the ERRαFL and ERRαΔNTD and contributed significantly in stability of the DNA bound ERRα proteins.

#### SASA analysis

Solvent-accessible surface area (SASA) was calculated on apo and DNA bound ERRαFL, ERRαΔNTD, ERRαDBD proteins (Fig. S1C). These data showed conformational flexibility and protein stability by quantifying the surface exposure to solvent molecules. Changes in SASA can reflect alterations in structural compactness upon LBD. The SASA profile showed that DNA bound ERRαΔNTD and ERRαDBD maintains stable solvent exposure throughout 100 ns simulation, with no significant deviation than apo proteins. In ERRαFL, minor inconsistency in surface area was observed till 30 ns simulation upon DNA binding and stabilized throughout 100 ns simulation. These data indicated that DNA binding does not induce increased solvent exposure to ERRα proteins unfolding or destabilization. This modest shift likely reflects local conformational adjustments required to accommodate DNA within the ERRα DNA binding pocket, which lead to minor changes in residue accessibility. Such changes are expected in ERRα proteins during DNA binding and do not indicate structural destabilization.

## Discussion

In present study, full length ERRα and its truncated constructs ERRαΔNTD and ERRαDBD domains were purified and characterized their interactions with the TMPRSS2-promoter DNA using biolayer interferometry technique. Secondary structures and thermal stability of the ERRα proteins were analysed using circular dichroism spectroscopy. Low resolution structures of the ERRαFL and ERRαFL-DNA complex were obtained using SAXS. Furthermore, dynamics simulation on apo and DNA-bound ERRα proteins were performed to investigate the conformational stability, structural dynamics and molecular mechanism underlying ERRα-DNA recognition.

The size-exclusion chromatography elution profiles, together with the standard Kav versus Log (Mr) calibration curve indicate that ERRαFL and ERRαDNTD proteins exist predominantly as dimers in solution. In contrast, the ERRαDBD exist in monomeric state. The circular dichroism analysis on apo and DNA bound ERRαFL, ERRαΔNTD, and ERRαDBD proteins showed native structures, and DNA binding induced only subtle changes in the secondary structure of ERRα proteins. Thermal denaturation analysis showed that DNA binding consistently increased the thermal stability of ERRα proteins. These data indicated that DNA binding stabilizes the folded conformations of the ERRα proteins by protein-DNA interactions and reducing conformational flexibility of the complex. These data were quite consistent to secondary structure analysis using CD, which showed minor variations in the secondary structural content in DNA bound ERRα proteins.

To investigate the effect of progressive domain deletion on the DNA-binding properties of full-length ERRα, N-terminal deleted ERRαΔNTD, and DNA-binding domain ERRαDBD were expressed and purified. The DNA-binding affinities of these proteins toward the ERRα response element (ERRE) were evaluated using biolayer interferometry. The isolated ERRαDBD binds weakly to the ERRE whereas ERRαFL and ERRαΔNTD binds substantially higher binding affinity then ERRαDBD. These results suggest that regions outside the isolated DNA-binding domain contribute to the efficient recognition and binding of ERRα to the ERRE.

Our data indicate that LBD, HR and DBD domains of ERRα may work together as a unified structural and functional unit, enabling efficient and high-affinity DNA binding. DNA binding to ERRα proteins stabilize the receptor dimerization and contributes to the overall conformation of the ERRα. In contrast, removing the N-terminal domain caused only a slight reduction in DNA-binding affinity, suggesting that the NTD contributes little to direct DNA recognition. ERRα-LBD domain is involved in the recruitment of co-regulators and transcriptional regulation, however contribute strongly in the stability of the protein-DNA complex and the facilitation of high-affinity DNA binding.

Small-angle X-ray scattering analysis showed that ERRαFL forms dimer in solution and fitted well with AlphaFold modelled ERRαFL homodimer. SAXS analysis of the ERRαFL-DNA complex further revealed a binding mode in which the dimeric receptor binds a 30 bp DNA duplex, two DNA-binding domains binding on the response element. These data provided low-resolution structural models of both apo and DNA-bound ERRαFL in solution, validated by overall structure predicted by AlphaFold3 server. It supports the mechanism in which dimerization of ERRα facilitates stable recognition of target DNA.

Structural models of apo and DNA bound ERRαFL, ERRαΔNTD, ERRαDBD complexes were subjected to 100 ns dynamics simulations to investigate their conformational stability and DNA-induced structural changes. Structural superposition and trajectory analyses revealed that the N-terminal domain (NTD) and hinge region (HR) are the most flexible regions of the receptor, whereas the core secondary structures of the ligand-binding domain (LBD) and DNA-binding domain (DBD) remained well preserved throughout the 100 ns dynamic simulation. The DNA duplex remained structurally stable in the ERRαFL-DNA and ERRαΔNTD-DNA complexes, while exhibited markedly greater conformational fluctuations in the ERRαDBD-DNA complex, consistent with its weaker DNA-binding affinity.

Sequence alignment and comparative structural analyses of ERRα with HNF4-α (PDB-4IQR) and Glucorticoid receptor (PDB-7PRW) revealed that residues involved in the LBD-LBD interface are not conserved. The organization of these nuclear receptors appears to be highly dependent on the constellation of non-conserved residues at the LBD-LBD interface, as well as the length and sequence composition of the hinge region. Moreover, ERRα residues involved in DNA recognition were partially conserved in the HNF-4α (PDB-4IQR) and Glucorticoid (PDB-7PRW) structures. Although LBD and DBD domains of ERRα are structurally conserved among HNF-4α and Glucorticoid receptors, the overall quaternary organization of these receptors diverges substantially.

## Conclusions

We have purified the ERRα proteins and characterized their secondary structures and thermal stability profile, low resolution structures and mechanism of recognition of the TMPRSS2 promoter DNA. Biophysical analysis showed that DNA binding stabilizes the folded conformation of ERRα proteins, likely by strengthening protein-DNA interactions and also showed modest changes in the secondary structures of ERRα proteins. DNA binding analysis showed that LBD and DBD modules of the ERRα are functionally integrated to establish the DNA binding affinity. The SAXS analysis revealed the low-resolution structure of the ERRαFL dimer and ERRαFL-TMPRSS2 DNA (2:1) complex. Dynamics simulation analysis revealed that the interdomain linkers of ERRα are intrinsically flexible, allowing DNA-binding domains (DBDs) and ligand-binding domains (LBDs) to undergo largely independent motions in apo and DNA-bound states. Upon DNA binding, the ERRα recognizes the DNA duplex, with the two LBDs forming a symmetric head-to-head dimer, while the two ERRαDBDs engage the left half and right half DNA recognition sites in a head-to-tail arrangement through sequence-specific contacts. The asymmetric organization of the ERRα-LBDs and ERRα-DBDs generates extensive interdomain interactions within the upstream protomer, accompanied by stabilization of a turn-like conformation in the linker region. These data showed that efficient DNA recognition by ERRα requires coordinated structural communication between the DBD and LBD, providing a mechanistic framework for understanding ERRα-mediated transcriptional regulation.

## Materials and Methods

### Expression and purification

Human Estrogen-related receptor alpha (ERRα, UniProt-P11474-1) plasmid was generated from cDNA derived from HT29 human colorectal cancer cells, which exhibit high endogenous ERRα expression. Total RNA was isolated and reverse-transcribed to obtain the full-length ERRα cDNA. The full length (ERRαFL, residues 1-423) and N-terminal deleted (ERRαΔNTD, residues 76-423) genes were amplified using Phusion high-fidelity DNA polymerase and cloned into *pET28a (+)* expression vector using *NheI* and *SalI* restriction sites. The DNA-binding domain of ERRα (ERRαDBD, residues 76–152) was cloned into *pET21c* (+) vector using *NheI* and *XhoI* restriction sites. All ERRα plasmids were verified by restriction-digestion.

The ERRαFL, ERRαΔNTD and ERRαDBD plasmids were transformed into *E. coli BL21(DE3)* cells. Single colony of ERRαFL and ERRαΔNTD was inoculated into LB media containing Kanamycin (50 µg ml^-1^), while ERRαDBD transformed colony was inoculated into LB media containing ampicillin (100 µg ml^-1^). Cells were cultured to mid-log phase (OD₆₀₀ ≈ 0.6), induced with 0.4 mM IPTG, and incubated at 18 °C for 16 h. The cells were harvested and lysed in buffer containing 20 mM Tris-HCl (pH 7.5), 500 mM NaCl, 10% glycerol, 8 mM β-mercaptethanol, 1 mM PMSF, 20 µg ml^-1^ lysozyme, and 2 µM ZnCl_2_. All proteins were purified using Ni-NTA affinity chromatography followed by size-exclusion chromatography on a Superdex 200(16/60) column equilibrated in 20 mM HEPES pH 7.5, 150 mM NaCl, 5% glycerol, and 8 mM β-mercaptoethanol. Protein purity were assessed by SDS-PAGE and measured the concentration using A280 measurement.

### DNA Preparation and complex formation with ERRα proteins

A 30-bp double-stranded DNA containing the Estrogen-related receptor response element (ERRE) sequence was prepared from synthetic oligonucleotides purchased from Sigms-Aldrich. The forward strand (5′-AGTCCAAGGTCGGTGGATCACAAGGTCAGG-3′) and its complementary reverse strand were dissolved in annealing buffer (20 mM Tris-HCl pH 7.5, 200 mM NaCl). The oligonucleotides were heated to 95 °C for 10 min and gradually cooled at room temperature for annealing. The annealed DNA was purified using QIAquick gel extraction columns (Qiagen). DNA purity was verified by agarose gel electrophoresis, and measured the concentration using absorbance at 260 nm.

For preparation of the ERRαFL-DNA complex, purified ERRαFL (∼ 0.26 mM) was incubated with pre-annealed double-stranded DNA (∼ 0.15 mM) at molar ratio (2: 1.2) in binding buffer containing 20 mM HEPES (pH 7.5), 100 mM NaCl, 1% (v/v) glycerol, 2 mM β-mercaptoethanol, and 10 µM MgCl_2_. The reaction mixture was incubated at 25 °C for 30 min to allow stable complex formation. An identical protocol was used for the preparation of the ERRαΔNTD-DNA and ERRαDBD-DNA complexes. The assembled ERRα-DNA complexes were subsequently used for biochemical, biophysical, and structural analysis experiments.

### Circular dichroism analysis

Circular dichroism (CD) spectroscopy was used to estimate the secondary structural contents of ERRαFL, ERRαΔNTD, ERRαDBD, and their respective DNA complexes. Far-UV CD spectra were recorded using a MOS-500 Spectrometer equipped with a 1 mm path-length quartz cuvette. All ERRα proteins (∼ 0.1-0.3 mg/ml) were prepared in 10 mM sodium phosphate buffer (pH 7.5), and spectra were collected over the wavelength range of 200-250 nm at 25 °C. Buffer spectra recorded under identical experimental conditions were subtracted from the corresponding protein spectra. Each CD spectra represents the average of three consecutive scans. Observed ellipticity was converted to mean residue ellipticity (MRE, deg·cm^2^·dmol^-1^·residue^-1^) and secondary structure content was estimated using the BeStSel server [21] using the following equation, [θ] *=* measured ellipticity (mdeg)/ (10 × c × l), where c is the protein concentration (μM), [θ] is the molar ellipticity in degree cm^2^ dmol^−1^, and l is the cuvette path length in mm.

Thermal denaturation analysis of the ERRαFL, ERRαΔNTD, ERRαDBD, and their respective DNA complexes was performed by CD spectroscopy. All ERRα proteins and their respective DNA complexes (∼ 0.1 mg mL^-1^) were placed in a 1 mm path-length quartz cuvette, and the ellipticity at 222 nm was recorded during thermal unfolding from 20 to 90 °C at 5 °C intervals. The melting temperature (T_m_) was determined by fitting the thermal unfolding curves using the Boltzmann sigmoidal equation in OriginPro software [22]. *Y*(*T*)=*A* 1+exp [*T*− *Tm/w*], where *Y(T)* is ellipticity at temperature T, *A* is the difference between folded and unfolded ellipticity, *Tm* is the melting temperature, and exp is the exponential function and *w* controls the width of the transition in the unfolding process,

### Bio-Layer Interferometry analysis

Bio-layer interferometry was performed using a Sartorius Octet R2 system to quantify the binding kinetics and affinities of ERRαFL, ERRαΔNTD and ERRαDBD to DNA. 5’-Biotin-labeled pre-annealed double stranded DNA (5’-Biotin-AGTCCAAGGTCGGTG GATCA CAAGGTCAGG-3’) and complementary oligonucleotide (3’-TCAGGTTCCAGCCA CCTAGTGTTCCAGTCC-5’) were pre-annealed and immobilized onto the streptavidin biosensors following sensor hydration in assay buffer (20 mM HEPES (pH 7.5), 200 mM NaCl, 10 µM MgCl_2_, and 2 mM β-mercaptoethanol) for 10 min. Immobilization was carried out for 200 s, followed by a baseline equilibration step for 300 s. Purified ERRαFL, ERRαΔNTD, and ERRαDBD proteins were prepared in BLI assay buffer and protein analytes were tested using a serial dilution series ranging from (2.5 µM to 0.078 µM), (3.0 µM to 0.093 µM) and (15 µM to 0.468 µM) respectively.

Association kinetics were measured by immersing DNA-loaded sensors into ERRα proteins solution for 200 s, followed by dissociation in assay buffer for 200 s. Reference sensors were subjected to identical association and dissociation steps in assay buffer without protein for background subtraction. Data were processed using Octet BLI analysis software [23]with reference subtraction applied prior to global fitting using a 2:1 binding model. Association rate constants (k_a_), dissociation rate constants (k_d_), and equilibrium dissociation constants (*K_D_*) were calculated from the fitted sensograms.

### Small-angle X-ray scattering analysis

Small-angle X-ray scattering data were collected using an Anton Paar SAXSpoint 2.0 instrument equipped with a two-dimensional EIGER R-series Hybrid Photon Counting (HPC) detector at the Indian Institute of Technology (IIT) Sonipat, India. X-rays wavelength (1.5418 Å) was used for SAXS data collection. 20 μL of ERRαFL (∼0.26 mM) and the ERRαFL-DNA complex (molar ratio of 2:1.2) were prepared in an equilibration buffer containing 20 mM HEPES (pH 7.5), 100 mM NaCl, 1% (v/v) glycerol, 2 mM β-mercaptoethanol and 10 µM MgCl_2_. The samples were loaded into a thermostated quartz capillary with a 1 mm path length and exposed to X-rays for 30 min. Three independent scattering frames were collected for each sample and the corresponding buffer, averaged, and used for subsequent analysis.

Scattering data were collected over a momentum transfer range of q = 0.02–20.7 nm^-1^, where q=4πsinθ/λ (2θ is the scattering angle and λ is the wavelength) and sample-to-detector distance of 1.075 m was used. Raw scattering data were processed using SAXSquant software (Anton Paar) for beam-centre correction, buffer subtraction, and desmearing of the scattering profiles. The resulting scattering intensity profiles [I(q) ∼ q] were analysed using the ATSAS software (version 3.0.1). The radius of gyration (Rg) was determined from Guinier analysis, while Kratky plots [I(q)q^2^ ∼ q] were used to assess the overall folding state and conformational flexibility of the ERRαFL and ERRαFL-DNA complex. The pair-distance distribution function P(r), and the maximum particle dimension (D_max_) were calculated by indirect fourier transformation using the GNOM program [24].The molecular weight (Mw) of the scattering particles was estimated using the SAXSMoW server [25].

Low-resolution *ab initio* molecular envelopes of ERRαFL and the ERRαFL-DNA complex were reconstructed using the DAMMIF, DAMMIN, and GASBOR programs within the ATSAS, Twenty independent DAMMIF models were generated for each sample and aligned and averaged using DAMAVER [26] to obtain a consensus envelope, which was subsequently refined using DAMMIN [27]. For the ERRαFL-DNA complex, rigid-body modelling was additionally performed using SASREF [28]. Theoretical scattering curves calculated from the structural models were fitted to the experimental SAXS profiles using CRYSOL [29] to evaluate the agreement between the models and the experimental data. The resulting SAXS envelopes and fitted atomic models were visualized using UCSF ChimeraX [30]. Details of SAXS data collection, data processing, and structural parameters are summarized in Table 1.

### Overall structure analysis of the ERRαFL and ERRαFL-DNA complex

Interface analysis of the dimeric ERRαFL and the ERRαFL-DNA (2:1) complex was performed using the PISA server [31], to identify the interfacial residues involved in homodimer formation. Protein-DNA interaction analysis of the ERRαFL-DNA complex, including identification of residues involved in DNA recognition and coordination of the Zn1 and Zn2 ions within the zinc-finger DNA-binding domain, was carried out using the DNAproDB server [32]. Structural superposition of ERRαFL and the ERRαFL-DNA complex was performed using the Superpose program [33] to evaluate the conformational differences associated with DNA binding.

### Sequence alignment and comparative structure analysis

The DALI server [34] analysis was performed on ERRαFL structure, which yielded closest structural homologs of the protein. Multiple sequence alignment was performed using the MultiAln [35] and ESPript [36] servers, aligning the ERRα sequence with its two closest structural homologs, HNF-4α-DR1 (PDB: 4IQR) and glucocorticoid receptor - DR-4 (PDB: 7PRW) complexes. The secondary structural elements of ERRα were mapped on the top of sequence alignment using ESPript program. Structural superposition of the ERRα-DNA complex with the HNF4α-DR-1 (PDB: 4IQR) and glucocorticoid receptor-DR-4 (PDB: 7PRW) complex structures was performed using the SUPERPOSE program.

### Molecular modelling and dynamics simulation analysis

Structural predictions were performed using the AlphaFold Server powered by AlphaFold 3 (AF3) [37]. The monomeric mode was used to predict the structures of all ERRα protein constructs, whereas the multimeric mode was used to model the complexes of ERRα proteins with 30-bp DNA. Five independent models were generated and model with highest confidence scores (pLDDT and pAE) were selected and analysed globally and at interfacial regions. Regions with pLDDT (predicted Local Distance Difference Test) values above 70 were considered reliable model. Overall model quality was further evaluated using multiple complementary structural validation metrics, including stereochemical assessment and structural consistency analyses. The PAE (Predicted Aligned Error) matrix was additionally examined to evaluate domain orientations and the confidence in inter-domain interactions. For multimeric models, inter-chain PAE values were analysed to assess the reliability of protein–protein interfaces and relative chain positioning. The full-length ERRα dimer structure was generated using crystal structure of the ERRα ligand-binding domain in complex with a PGC-1α peptide (PDB −1XB7) and ERRαFL-DNA complex using structure of HNF-4α-DR1 (PDB-4IQR) as the templates. The hinge region connecting the DNA-binding and ligand-binding domains was refined by loop modelling. Further analysis of complex properties, such as surface area and binding energies, was performed using the PDBePISA server. PDBsum [38] was employed to analyse the interface residues of the complexes, providing insights into key interacting amino acids.

Molecular dynamics (MD) simulation was performed to evaluate the conformational stability and protein-DNA interactions of ERRαFL, ERRαΔNTD, and ERRαDBD in complex with DNA (5’-AGTCCAAGGTCGGTGGATCACAAGGTCAGG-3’). All simulations were carried out using GROMACS (versions 2022.3) [39] program using the AMBER99SB-ILDN force field [40] for protein and AMBER94 parameters for nucleic acid. Systems were solvated in a cubic box with TIP3P water molecules [41] maintaining a minimum distance of 10 Å between the solute and box edges and neutralized by the addition of Na⁺ and Cl⁻ ions. Energy minimization was performed using 50,000 steps of the steepest descent algorithm. Systems were equilibrated in two phases, constant volume and temperature (NVT) for 100 ps, followed by constant pressure and temperature (NPT) for 100 ps at 300 K and 1 bar. Production MD simulations were conducted under NPT conditions for 100 ns using the V-rescale thermostat and Berendsen barostat, with a 2-fs integration time steps and coordinates were saved every 10 ps. LINCS algorithm was used to constrain all bonds, Particle Mesh Ewald (PME) method for long-range electrostatic interactions. The pre and post simulation, the structures were superposed to examine the protein stability, conformational changes and protein-protein interface area. XMGRACE [42] was used to plot the root-mean-square deviation (RMSD), root-mean-square fluctuation (RMSF), Rg, hydrogen bonds, solvent accessible surface area and PyMOL [43] for visualization.

## Authors contributions

**K Chandrashekhara:** Conceptualization, experimental design, methodology, investigation, software, data curation, formal analysis, validation, visualization, carried out all experiments, wrote and edited the manuscript.

**AKS:** Conceptualization, methodology, Writing-Original Draft Preparation, Writing and Editing, Supervision, Resources, Project Administration, Fund generation.

## Acknowledgements

Chandrashekhara gratefully acknowledges the Department of Biotechnology (DBT) for providing fellowship for Ph.D. research. The authors sincerely thank Ms. Aditi Rattan for her support in generating the MD simulation data. The authors acknowledge the BRAHM: High-Performance Computational facility of the Indian Biological Data Centre, Regional Centre for Biotechnology, Faridabad, INDIA (https://ibdc.rcb.res.in/; DBT Grant no. BT/TCB/IBDC/2019). The authors gratefully acknowledge financial support from the Department of Biotechnology (DBT) through grant BT/PR45101/DRUG/ 134/121/2022 and from the Indian Council of Medical Research (ICMR). The authors also acknowledge the support of the DBT-BUILDER programme (Grant No. BT/INF/22/ SP45382/2022) for establishing the Central Instrumentation Facility (CIF) at Jawaharlal Nehru University (JNU) and for providing institutional funding. The authors further acknowledge the Central Instrumentation Facility (CIF), SLS), JNU, they also thank the Central Research Facility (CRF), IIT Delhi, for providing access to SAXS and CD spectroscopy facilities.

## Conflict of interests

The authors declare that they have no conflict of interest.

## Data availability statements

Data will be made available on request.

## Abbreviations

ERRα: Estrogen related receptor Alpha
ERRα proteins: (ERRαFL, ERRαΔNTD, ERRαDBD)
ERRαFL: Full length ERRα
ERRαΔNTD: NTD deleted ERRα
ERRαDBD: DNA binding domain of ERRα
DNA: TMPRSS2 promoter DNA
TMPRSS2: Transmembrane serine protease-2
PDB: Protein Data Bank
CD: Circular Dichroism
SAXS: small angle X-ray scattering
BLI: Bio-layer interferometry
Ni-NTA: Nickel-Nitriloacetic Acid
SDS PAGE: Sodium Dodecyl Sulphate Polyacrylamide Gel Electrophoresis
MSA: Multiple Sequence Alignment
MD: Molecular dynamics
RMSD: Root Mean Square Deviation
RMSF: Root mean Square Fluctuation
Rg: Radius of Gyration
SASA: Solvent Accessible Surface Area

---

**Fig. S1.**

(A) Solvent-accessible surface area (SASA) plot, illustrating alterations in protein-solvent interactions for ERRαFL and ERRαFL-DNA complex. The lower panels show PDF distributions, providing further insights into the stability of the systems. (B) The time evolution of intramolecular hydrogen bonds within ERRαFL and their stability in apo and DNA-bound states. (C) The time evolution of Intermolecular hydrogen bonding between ERRαFL and ERRαFL-DNA complex.

## Notes

### Competing Interest Statement

The authors have declared no competing interest.

