## Supplementary figures and images for "Structural and biochemical analysis of the Estrogen-Related Receptor α and complex with TMPRSS2 promoter DNA"

### fig.s1

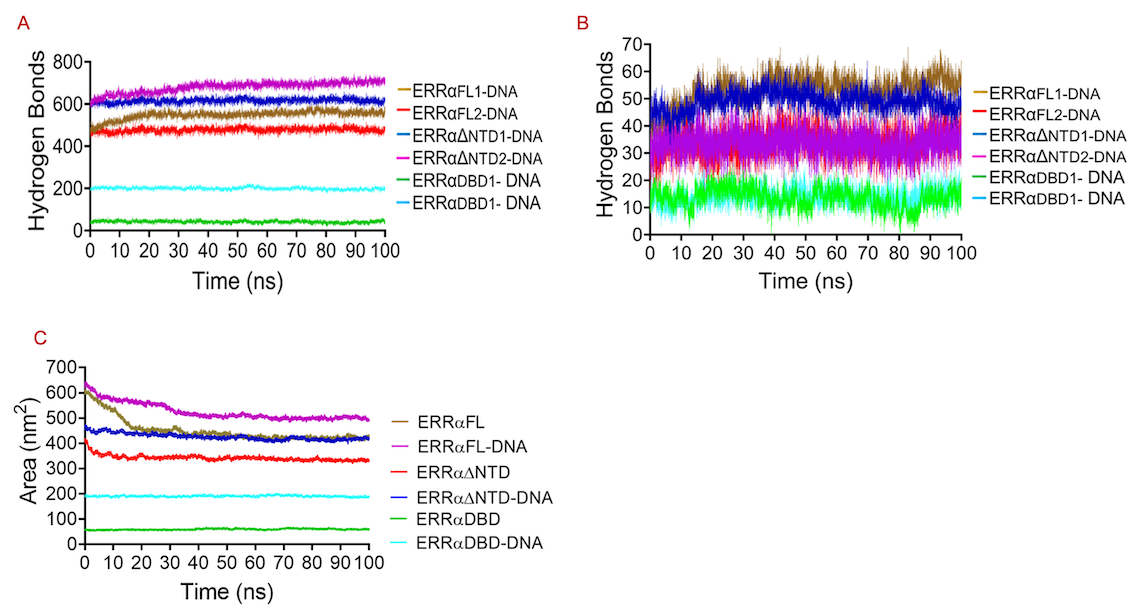

### fig.s2

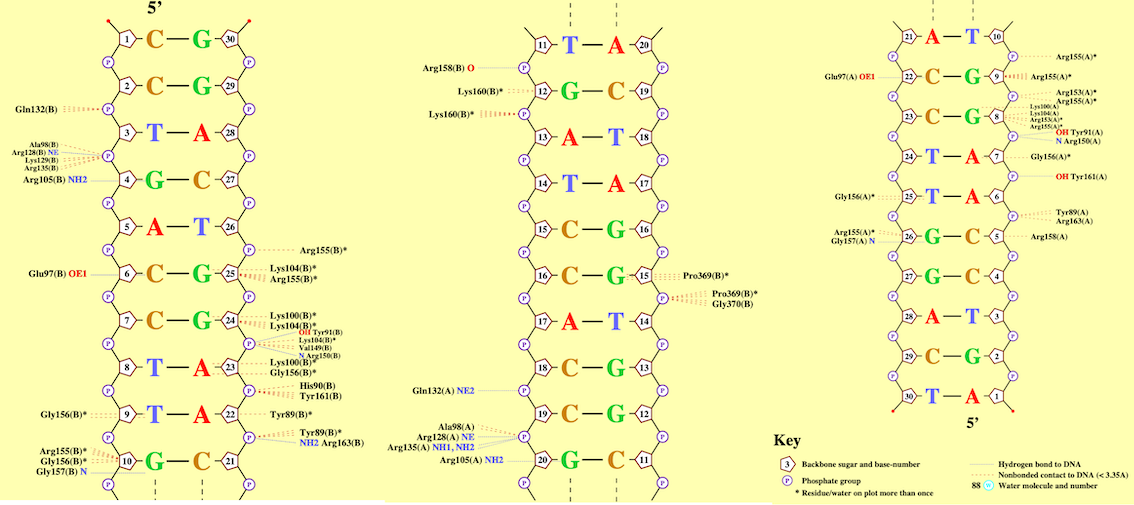
