## Supplementary material for "Structural and biochemical analysis of the Estrogen-Related Receptor α and complex with TMPRSS2 promoter DNA": Table s1

Table S1. Hydrogen bonds and salt bridges between ERRαFL monomers

Hydrogen bonds

| | ATOM 1 | | | | | | ATOM 2 | | | | | |
|---|---|---|---|---|---|---|---|---|---|---|---|---|
| | Atom no. | Atom name | Res name | Res no. | Chain | | Atom no. | Atom name | Res name | Res no. | Chain | Distance |
| 1. | 2310 | NE2 | GLN | 311 | A | <--> | 5719 | O | ASP | 338 | B | 2.34 |
| 2. | 2346 | NH1 | ARG | 315 | A | <--> | 5722 | OD1 | ASP | 338 | B | 2.40 |
| 3. | 2347 | NH2 | ARG | 315 | A | <--> | 5708 | OD1 | ASN | 336 | B | 2.51 |
| 4. | 2516 | OD1 | ASN | 336 | A | <--> | 5539 | NH2 | ARG | 315 | B | 2.53 |
| 5. | 2517 | ND2 | ASN | 336 | A | <--> | 6061 | O | LEU | 383 | B | 3.08 |
| 6. | 2527 | O | ASP | 338 | A | <--> | 5502 | NE2 | GLN | 311 | B | 2.35 |
| 7. | 2530 | OD1 | ASP | 338 | A | <--> | 5538 | NH1 | ARG | 315 | B | 2.42 |
| 8. | 2651 | OE2 | GLU | 353 | A | <--> | 6038 | NE | ARG | 380 | B | 2.73 |
| 9. | 2700 | O | LEU | 360 | A | <--> | 5999 | NH2 | ARG | 375 | B | 2.74 |
| 10. | 2807 | NH2 | ARG | 375 | A | <--> | 5892 | O | LEU | 360 | B | 2.74 |
| 11. | 2846 | NE | ARG | 380 | A | <--> | 5843 | OE2 | GLU | 353 | B | 2.12 |
| 12. | 2869 | O | LEU | 383 | A | <--> | 5709 | ND2 | ASN | 336 | B | 3.08 |

Salt bridges

| | ATOM 1 | | | | | | ATOM 2 | | | | | |
|---|---|---|---|---|---|---|---|---|---|---|---|---|
| | Atom no. | Atom name | Res name | Res no. | Chain | | Atom no. | Atom name | Res name | Res no. | Chain | Distance |
| 1. | 2346 | NH1 | ARG | 315 | A | <--> | 5722 | OD1 | ASP | 338 | B | 2.40 |
| 2. | 2530 | OD1 | ASP | 338 | A | <--> | 5538 | NH1 | ARG | 315 | B | 2.42 |
| 3. | 2650 | OE1 | GLU | 353 | A | <--> | 6041 | NH2 | ARG | 380 | B | 2.27 |
| 4. | 2682 | OE1 | GLU | 357 | A | <--> | 5998 | NH1 | ARG | 375 | B | 3.54 |
| 5. | 2683 | OE2 | GLU | 357 | A | <--> | 6007 | NE | ARG | 376 | B | 3.59 |
| 6. | 2806 | NH1 | ARG | 375 | A | <--> | 5874 | OE1 | GLU | 357 | B | 3.57 |
| 7. | 2815 | NE | ARG | 376 | A | <--> | 5875 | OE2 | GLU | 357 | B | 3.63 |
| 8. | 2846 | NE | ARG | 380 | A | <--> | 5842 | OE1 | GLU | 353 | B | 2.12 |
