## Supplementary material for "Structural and biochemical analysis of the Estrogen-Related Receptor α and complex with TMPRSS2 promoter DNA": Table s2

Table S2. Hydrogen bonds and salt bridges between ERRαFL-DNA Complex

Hydrogen bonds

| | Atom 1: Atom no. | Atom 1: Atom name | Atom 1: Res name | Atom 1: Res no. | Atom 1: Chain | | Atom 2: Atom no. | Atom 2: Atom name | Atom 2: Res name | Atom 2: Res no. | Atom 2: Chain | Distance |
|---|---|---|---|---|---|---|---|---|---|---|---|---|
| 1. | 2346 | NH1 | ARG | 315 | A | <--> | 5721 | OD1 | ASP | 338 | B | 2.13 |
| 2. | 2347 | NH2 | ARG | 315 | A | <--> | 5708 | OD1 | ASN | 336 | B | 2.67 |
| 3. | 2517 | OD1 | ASN | 336 | A | <--> | 5538 | NH2 | ARG | 315 | B | 2.72 |
| 4. | 2516 | ND2 | ASN | 336 | A | <--> | 6060 | O | LEU | 383 | B | 3.06 |
| 5. | 2530 | OD1 | ASP | 338 | A | <--> | 5537 | NH1 | ARG | 315 | B | 2.17 |
| 6. | 2614 | OE2 | GLU | 349 | A | <--> | 6040 | NH2 | ARG | 380 | B | 2.44 |
| 7. | 2683 | OE2 | GLU | 357 | A | <--> | 5998 | NH2 | ARG | 375 | B | 2.60 |
| 8. | 2807 | NH2 | ARG | 375 | A | <--> | 5874 | OE2 | GLU | 357 | B | 2.54 |
| 9. | 2849 | NH2 | ARG | 380 | A | <--> | 5805 | OE2 | GLU | 349 | B | 2.49 |
| 10. | 2869 | O | LEU | 383 | A | <--> | 5707 | ND2 | ASN | 336 | B | 3.03 |

Salt bridges

| | Atom 1: Atom no. | Atom 1: Atom name | Atom 1: Res name | Atom 1: Res no. | Atom 1: Chain | | Atom 2: Atom no. | Atom 2: Atom name | Atom 2: Res name | Atom 2: Res no. | Atom 2: Chain | Distance |
|---|---|---|---|---|---|---|---|---|---|---|---|---|
| 1. | 2346 | NH1 | ARG | 315 | A | <--> | 5721 | OD1 | ASP | 338 | B | 2.13 |
| 2. | 2530 | OD1 | ASP | 338 | A | <--> | 5537 | NH1 | ARG | 315 | B | 2.17 |
| 3. | 2613 | OE1 | GLU | 349 | A | <--> | 6040 | NH2 | ARG | 380 | B | 2.44 |
| 4. | 2651 | OE2 | GLU | 353 | A | <--> | 6008 | NH1 | ARG | 376 | B | 3.01 |
| 5. | 2650 | OE1 | GLU | 353 | A | <--> | 6037 | NE | ARG | 380 | B | 2.80 |
| 6. | 2682 | OE1 | GLU | 357 | A | <--> | 5998 | NH2 | ARG | 375 | B | 2.60 |
| 7. | 2683 | OE2 | GLU | 357 | A | <--> | 6006 | NE | ARG | 376 | B | 3.51 |
| 8. | 2712 | OE1 | GLU | 361 | A | <--> | 5998 | NH2 | ARG | 375 | B | 3.99 |
| 9. | 2807 | NH2 | ARG | 375 | A | <--> | 5873 | OE1 | GLU | 357 | B | 2.54 |
| 10. | 2807 | NH2 | ARG | 375 | A | <--> | 5903 | OE1 | GLU | 361 | B | 3.89 |
| 11. | 2817 | NH1 | ARG | 376 | A | <--> | 5842 | OE2 | GLU | 353 | B | 2.93 |
| 12. | 2815 | NE | ARG | 376 | A | <--> | 5874 | OE2 | GLU | 357 | B | 3.47 |
| 13. | 2849 | NH2 | ARG | 380 | A | <--> | 5804 | OE1 | GLU | 349 | B | 2.49 |
| 14. | 2846 | NE | ARG | 380 | A | <--> | 5841 | OE1 | GLU | 353 | B | 2.85 |
